# L1CAM signaling through planar cell polarity generates SOX2^+^ metastatic progenitors in lung adenocarcinoma

**DOI:** 10.1101/2025.08.22.671773

**Authors:** Jin Suk Park, Yasemin Kaygusuz, Carson Kenum, Lan He, Mohamed I. Gatie, Xueqian Zhuang, Roshan Sharma, Ronan Chaligné, Umesh K. Bhanot, Richard P. Koche, Tuomas Tammela, Anna-Katerina Hadjantonakis, Charles M. Rudin, Joan Massagué

## Abstract

Plasticity transitions during carcinoma progression generate fetal-like progenitor states with metastatic capacity. How these progenitors emerge and persist during tumor progression remains unknown. Here, we elucidate a process that drives the emergence of SOX2^+^ metastatic progenitors in lung adenocarcinomas (LUAD). LUAD cells in the tumor invasive front and distant metastases express the cell adhesion molecule L1CAM, a marker of regenerative epithelial progenitors and a mediator of cell-basement membrane and cell-cell interactions. L1CAM-mediated adhesion to perivascular basement membrane is known to stimulate the proliferation of extravasated micrometastatic cells. We now identify a distinct and broader role of L1CAM as promoter of the SOX2+ LUAD progenitor state. We show that L1CAM at cell-cell interfaces promotes the assembly of the planar cell polarity (PCP) complex in metastatic LUAD progenitors. L1CAM-dependent PCP acting through a non-canonical WNT signaling activates c-Jun, which cooperates with the chromatin remodeling factor CHD1 to drive *SOX2* expression and metastatic activity. This axis sustains the tumor-initiating and regenerative capacity of LUAD progenitor cells. By illuminating the role of L1CAM and PCP signaling in the generation of SOX2^+^ LUAD progenitors, our findings identify potential new targets to treat metastatic cancer.

## Introduction

Metastasis, the principal cause of cancer-related mortality, entails a complex cascade involving the dissemination, dormancy, adaptation, and outgrowth of malignant cells at distant organ sites^1^. While numerous molecular and cellular mechanisms have been implicated in organ-specific patterns of metastasis across diverse cancer types, a unifying requirement is the capacity of disseminated tumor cells to reinitiate growth despite multiple environmental and therapeutic barriers. In epithelial cancers, this regenerative capacity is often acquired through the adoption of phenotypic states, typically driven by non-genetic mechanisms during tumor progression and therapeutic stress^2–4^. A growing body of evidence suggests that phenotypic plasticity underlies the emergence of fetal-like progenitor phenotypes with metastatic capacity. In lung^5–8^, colorectal^9–13^, and prostate adenocarcinomas^14–16^, tumors originating from adult homeostatic stem cells progressively acquire multilineage plasticity, giving rise to distinct progenitor states that express developmental programs associated with metastatic competence. However, the mechanisms governing the emergence and maintenance of these progenitor phenotypes throughout tumor progression remain poorly understood.

Lung adenocarcinoma (LUAD), a leading cause of cancer death worldwide, exemplifies these dynamic lineage transitions. LUAD originates from alveolar type II (AT2) cells with stem-like features^5, 17, 18^, which progressively lose their lineage identity and give rise to heterogeneous populations expressing developmental progenitor markers^6^ and exhibiting features of multilineage plasticity^5, 7^. LUAD metastases recapitulate a continuum of progenitor stages reflective of fetal lung development^6^, ranging from anterior endoderm-like SOX2⁺ cells to more differentiated bronchoalveolar progenitor phenotypes. Notably, SOX2⁺ cells derived from primary LUAD tumors display hallmark properties of metastasis-initiating cells, including heightened metastatic seeding capacity, the ability to enter and exit dormancy, immune evasion, and aggressive multi-organ colonization upon immune escape^19, 20^. Yet, the regulatory mechanisms sustaining SOX2 expression and progenitor identity during LUAD progression and metastasis remain elusive.

L1 cell adhesion molecule (L1CAM) is implicated as a key regulator of regenerative progenitors during injury repair and cancer metastasis. L1CAM is a transmembrane protein that binds laminins for cell spreading on basement membranes and alternatively forms homophilic L1CAM-L1CAM interactions at cell-cell interfaces^21, 22^. Normally confined to the central nervous system and distal renal tubules, L1CAM expression is induced upon injury and disruption of adherens junctions in epithelial tissues^9^. L1CAM is frequently expressed in solid tumors including LUAD^23^, and is correlated with poor prognosis^24, 25^. L1CAM promotes metastatic colonization across diverse types of carcinomas in experimental models^9, 26, 27^. It mediates the spreading of extravasated cancer cells on perivascular basement membranes, inducing mechanotransduction through β1 integrin signaling and YAP and MRTF activation^27^. Although this role of L1CAM is important for the initial proliferation of extravasated cancer cells^27^, L1CAM in tumors is mainly localized at cancer cell-cell interfaces where homophilic L1CAM binding interactions can take place^9, 26, 27^. This suggests a broader role for L1CAM in the emergence of metastasis-competent progenitor phenotypes. Indeed, recent findings in colorectal cancer have linked L1CAM to the transition toward a high-plasticity, fetal-like progenitor state capable of adapting to metastatic microenvironments^10^.

In this study, we investigate the role of L1CAM as a critical determinant of metastatic progenitor identity in LUAD. Using human LUAD samples, a genetically engineered mouse model, and tumor organoids derived from both sources, we demonstrate that L1CAM is dispensable for the initiation of primary lung tumors but important for the acquisition of metastatic competence. We found that LUAD primitive SOX2^+^ progenitors exhibit peak expression of L1CAM accompanied with peak expression of the planar cell polarity (PCP) complex and promotes the assembly of this complex in SOX2^+^ cells. PCP is a key regulator of epithelial morphogenesis that triggers non-canonical WNT signaling via Frizzled (FZD)-driven activation of Jun N-terminal kinase (JNK)^28–31^. While PCP is fundamental to morphogenesis, its role in tumorigenesis remains largely unknown^31^. We show that L1CAM promotes PCP complex assembly, initiating a signaling cascade that activates *SOX2* transcription for metastatic competence in LUAD progenitors.

## Results

### Emergence of L1CAM^+^ cells during LUAD progression

Although L1CAM expression has been noted in several cancers^24^, there is limited information on L1CAM expression in LUAD^23^. To bridge this gap, we profiled the distribution of L1CAM^+^ cells in primary and metastatic tumor samples from LUAD patients. L1CAM^+^ cells were not detected by immunohistochemical staining (IHC) in normal human lung epithelium and were rare in primary tumors but were prevalent in malignant pleural effusions and metastases across various organs (Fig. 1a,b and Extended Data Fig. 1a,b). We also profiled a collection of 106 patient-derived xenograft models (PDXs) established from human primary or metastatic LUAD tumors by our group. L1CAM^+^ cells were abundant in PDXs established from primary tumors or metastases (Fig. 1c and Extended Data Fig. 1a,b), suggesting that the regrowth of primary tumors as PDXs enriched for L1CAM^+^ cells. L1CAM^+^ cells were also present in PDX-derived spheres grown under organoid culture conditions (referred to as PDX tumoroids), as determined by L1CAM immunofluorescence (IF) staining (Extended Data Fig. 1c-e).

**Figure 1.**
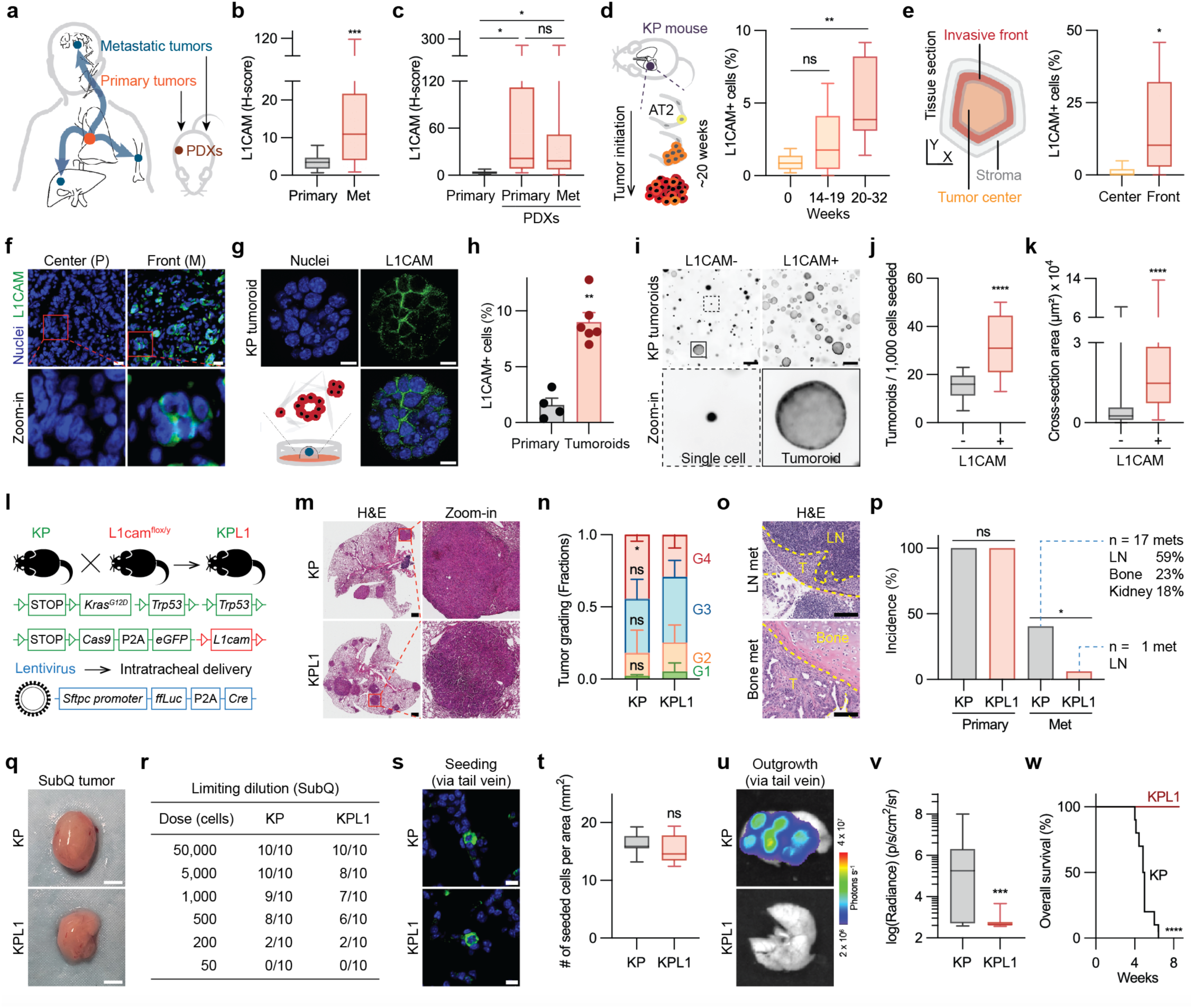
Emergence of L1CAM^+^ metastatic cells during LUAD progression. **a**, Schematic of metastatic LUAD and patient-derived xenograft (PDX) engraftment. **b**, Box and whisker plots of L1CAM IHC H-score in patient-derived primary tumor and metastasis samples. Primary tumor, *n =* 15; metastases, *n =* 54. \*\*\**P* = 0.0002. **c**, Box and whisker plots of L1CAM IHC H-score in the primary tumors in (**b**) and in PDXs derived from primary tumor or metastases. Primary tumors, *n =* 15; primary tumor-derived PDXs, *n =* 36; metastasis-derived PDXs, *n =* 70. ns, *P* = 0.8811; \**P* = 0.0294 (left), 0.0420 (right). **d**, Schematic of the oncogenic transformation of AT2 cells into lung adenocarcinoma in the KP GEMM^32^ (*left panel*), and box and whisker plots of the percentage of L1CAM^+^ cells in the KP mouse lungs and tumor areas at different time points after lentivirus instillation (*right panel*). Week 0, *n =* 6; week 14-19, *n =* 6; week 20-32, *n =* 15. ns, *P* = 0.5710; \*\**P* = 0.0032. **e**, Schematic of the tumor tissue section (*left panel*), and box and whisker plots of the percentage of L1CAM^+^ cells in the tumor center and invasive front (*right panel*). Tumor center, *n* = 300 cells; invasive front, *n* = 258 cells. *N* = 7 image frames. \**P* = 0.0152. **f**, L1CAM immunofluorescence (IF) staining of LUAD patient tissue samples at the tumor center (P, papillary) or the invasive front (M, micropapillary). Magnified regions are indicated in red boxes. Scale bar, 20 μm. **g**, Confocal microscopy image of L1CAM IF staining and Hoechst counterstaining of nuclei in KP tumoroids grown for 5 days. Schematic of tumoroids generated from KP-derived cancer cells (*lower right panel*). Scale bar, 10 μm. **h**, Percentage of L1CAM^+^ cells in KP primary tumors versus tumoroids. Primary tumor, *n =* 4 experiments; tumoroids, *n =* 6 experiments. Mean ± s.e.m. \*\**P* = 0.0095. **i**, Widefield fluorescence microscopy image of KP tumoroids stained with calcein AM in KP tumor cells sorted by L1CAM expression and grown as tumoroids for 7 days. The magnified region showing a single cell is indicated by a dotted box and that of a tumoroid by a solid box. Scale bar, 400 μm. **j**, Box and whisker plots of tumoroids formed per 1,000 cells after 7 days in culture. L1CAM^−^, *n =* 16; L1CAM^+^, *n =* 16. \*\*\*\**P* < 0.0001. **k**, Box and whisker plots of cross-section area per tumoroid in the experiment of panel (**I**). L1CAM^-^, *n =* 807; L1CAM^+^, *n =* 477. \*\*\*\**P* < 0.0001. **l**, Schematic representation of the generation of *L1cam* knockout KP LUAD GEMMs. **m**, H&E staining of KP and KPL1 primary tumors (26 week post-Cre) followed by their histopathological grading using an automated deep neural network. Magnified regions are shown in a red square. Scale bar, 1 mm. **n**, Fraction of KP and KPL1 tumors (22-26 week post-Cre) based on histopathological grading. Grade 1, *green*; Grade 2, *orange*; Grade 3, *blue*; Grade 4, *red*. KP, *n =* 4; KPL1, *n* = 4. Mean ± S.D. **o**, H&E staining of KP metastasis in the subcapsular sinus of the lymph node (top panel) or ribcage bone (bottom panel). Invasive front is shown with a dotted yellow line. T, tumor; LN, lymph node. Scale bar, 100 μm. **p**, Incidence of primary tumor and spontaneous metastases upon viral transduction in KP and KPL1 mice. The organ specificity of spontaneous metastasis is shown as a percentage of the total metastatic tumors. LN, lymph node. KP primary, *n* = 46; KPL1 primary, *n* = 30; KP met, *n* = 42; KPL1 met, *n* = 16. ns, *P* = 1; \**P* = 0.0122. **q**, Representative images of subcutaneous tumors formed 6 weeks after inoculating 500 KP or KPL1 cells in athymic mice. Scale bar, 2 mm. **r**, Limiting dilution assay of subcutaneous tumor formation at various doses of KP or KPL1 cells in athymic mice. *n =* 10 for each condition. **s**, Fluorescent images of GFP^+^ KP or KPL1 cells seeded in lungs at week 1 after tail vein injection (10^5^ cells each) into athymic mice. Scale bar, 10 μm. **t**, Quantification of KP and KPL1 cells seeded in lungs from the experiment in (**S**). KP, *n* = 9; KPL1, *n* = 10. ns, *P* = 0.3154. **u**, Representative image of *ex vivo* lung BLI at week 5 after tail vein injection of single-cell suspension of KP and KPL1 tumoroids (2 x 10^4^ cells) into athymic mice. **v**, Quantification of *ex vivo* lung BLI signal in the experiment in (**u**). KP, *n =* 28; KPL1, *n =* 14. \*\*\**P* = 0.0006. **w**, The KM plot showing the overall survival after performing tail vein injections with KP or KPL1 cells. KP, *n* = 10; KPL1, *n* = 10. \*\*\*\**P* < 0.0001. Statistical significance was assessed using the two-tailed Mann-Whitney test (**b**,**e**,**h**,**j**,**k**,**n**,**t**,**v**), one-way analysis of variance followed by the Tukey test (**c**,**d**), two-tailed Fisher’s exact test (**p**) or log-rank (Mantel-Cox) test (**o**,**w**). Data are shown as a box (median ± 25-75%) and whisker (maximum to minimum values) plot (**b**-**e**,**j**,**k**,**t**,**v**).

To investigate the emergence of L1CAM^+^ cells during LUAD progression we used the genetically engineered mouse model (GEMM) *Kras*^LSL-G12D/+^;*Trp53*^flox/flox^;*Rosa26*^LSL-Cas9-EGFP^ (KP model) in which intratracheal instillation of Lenti-SPC-*ffLuc*-P2A-*Cre* lentivirus initiates formation of lung tumors that progress to adenocarcinoma over a 5-month period (Extended Data Fig. 2a)^32^. L1CAM^+^ cells were absent in normal mouse lungs but became evident at the invasive front as the tumors developed (Fig. 1d,e and Extended Data Fig. 2b), possibly reflecting L1CAM expression upon disruption of cancer cell junctions^9^. The L1CAM^+^ cells were concentrated in regions of micropapillary histology, which is associated with LUAD recurrence and metastasis^33^, but not in lepidic, acinar, papillary, or solid histology regions (Fig. 1f and Extended Data Fig. 2c-e). Notably, L1CAM^+^ cells were enriched in autochthonous metastases to mediastinal lymph nodes compared to primary tumors (Extended Data Fig. 2f,g), mirroring our observations in patient-derived primary tumors versus metastases. KP tumoroids grown from primary tumors (Extended Data Fig. 2h-j) also exhibited an enrichment for L1CAM^+^ cells compared to the source tumors (Fig. 1g,h). Of note, in human and mouse LUAD tissue samples (Fig. 1f and Extended Data Fig. 1a,2a,j) as well as in tumoroids (Fig. 1g and Extended Data Fig. 1d,e), L1CAM was predominantly localized at cell-cell interfaces. L1CAM^high^ cancer cells sorted from KP lung tumors showed an elevated capacity for growth as tumoroids compared to L1CAM^low^ cells (Fig. 1i-k).

### L1CAM promotes LUAD metastasis

To assess the role of L1CAM in LUAD tumor growth and metastasis, we generated a *L1cam*-null KP GEMM (referred to as the KPL1 model) by crossing KP mice with mice harboring a conditional *L1cam* null allele (*L1cam*^flox/y^)^34^ (Fig. 1l). Tumor formation in both KP and KPL1 mice progressed at similar rates, as determined by methods including quantitative bioluminescence (BLI) of a vector-encoded firefly luciferase, micro-computed tomography (micro-CT) scanning, and histologic analysis of tumor tissue sections (Extended Data Fig. 3a-e). KP and KPL1 lung tumors showed a similar histology, featuring carcinoma cells with polygonal morphology, a mix of tubular and solid growth patterns, and extensive invasion of the adjacent lung parenchyma (Fig. 1m). While most tumors were of intermediate grade overall, histopathological grading using an automated deep neural network analysis showed a slightly greater proportion of grade-4 regions in KP tumors compared to time- and size-matched KPL1 tumors (Fig. 1m,n).

We analyzed the presence of autochthonous metastases in KP and KPL1 mice as their primary tumor burden reached a total mouse chest BLI flux of 10^7^ photons/sec. KP and KPL1 mice reached this target tumor burden with the same kinetics (Extended Data Fig. 3f), which was the burden chosen for tumor harvesting to generate tumoroids. Despite KP and KPL1 mice displaying similar tumor growth rate, burden, and histopathological grade, the post-mortem examination of mice reaching this target burden at the same time revealed the presence of metastases to lymph nodes, adrenal glands, kidneys, and/or ribcage bones in 17/42 KP mice but only in 1/16 KPL1 mice (Fig. 1o,p). These results agreed with our previous findings in colorectal cancer^9^ and suggested a limited contribution of L1CAM to carcinoma initiation but a critical role of L1CAM in metastatic progression in LUAD.

To further assess the role of L1CAM in LUAD metastasis, we sorted GFP^+^ cancer cells from mouse lung tumors and grew them as tumoroids in vitro or transduced them with a lentiviral thymidine kinase-GFP-luciferase fusion protein vector prior to subcutaneous inoculation into the flanks of mice for tumor formation activity analysis. KP tumor cell suspensions, which predominantly consist of L1CAM^−^ cells, and KPL1 cell suspensions were comparable in their capacity to grow as tumoroids (Extended Data Fig. 3g) or as subcutaneous tumors (Fig. 1q,r). Tumoroid cell suspensions were inoculated into athymic nude mice through three different routes: via the tail vein for metastatic colonization in the lungs—a major site of LUAD metastasis; intratracheally to study tumor spread through air spaces, a clinically relevant mode of LUAD intrapulmonary metastasis^35, 36^; and via the left cardiac ventricle into the arterial circulation to assess multi-organ metastasis (Extended Data Fig. 3h). The metastatic seeding potential of KP or KPL1 cells was similar, as determined by quantification of GFP^+^ cells in the lungs one week after intravenous inoculation (Fig. 1s,t). KP cells generated lung metastases via these three routes and metastasis to kidney, brain, and liver via intracardiac inoculation, whereas KPL1 cells failed to form metastasis via all three routes (Fig. 1u-w and Extended Data Fig. 3i-r).

To determine the role of L1CAM in the metastatic activity of LUAD PDXs, we knocked down L1CAM levels in two different PDX models (Ru631 and Ru323) using two independent *L1CAM* shRNAs in each model. Injection of PDX cell suspensions into the tail vein of immunodeficient *Foxn1^nu^*athymic nude mice showed a decrease in metastatic activity which was proportional to the extent of L1CAM depletion (Extended Data Fig. 3s-x). Collectively, these results suggested that L1CAM is not essential for the initiation of primary LUAD tumors but is critical for metastasis by disseminated LUAD cells.

### SOX2^+^ LUAD cells act as metastatic progenitors

We explored the relationship between L1CAM and the lung developmental stages previously noted in human LUAD metastases^6^. Key transcription factors defining the successive progenitor stages during lung development include the pluripotency factor SOX2, characteristic of stem cells and endodermal and neural progenitors, the homeobox factor NKX2-1 (NK2 homeobox 1), which marks early lung development, the pioneer factor FOXA2 (forkhead box A2) which specifies intermediate lung progenitors, and HMGA2 (high motility group A2) and SOX9, which mark late-stage distal alveolar structures^37–40^. To characterize with more granularity the spatiotemporal emergence of SOX2^+^, NKX2-1^+^ and SOX9^+^ progenitors during lung development, we used IF staining on iDISCO-cleared whole mouse embryos across sequential embryonic stages (Fig. 2a, b and Extended Data Fig. 4a,b). At embryonic day (E)8.5, preceding the emergence of the lung bud primordium within the anterior foregut, cells of the anterior foregut were SOX2^+^. By E10.5 when the lung buds have ventrally emerged and initiated branching and the trachea is separating from the esophagus, there was a distinct separation of cells into SOX2^+^ esophagus, NKX2-1^+^ trachea and lung buds, and SOX9^+^ cells in the distal tips of the branching lung buds which contain alveolar progenitors. Thus, cells that eventually express SOX9 will have previously expressed NKX2.1 and SOX2. These results refine the previously delineated sequential expression of SOX2, NKX2-1, and SOX9 during lung development^37–40^, underscoring the role of these factors in the specification of lung progenitors.

**Figure 2.**
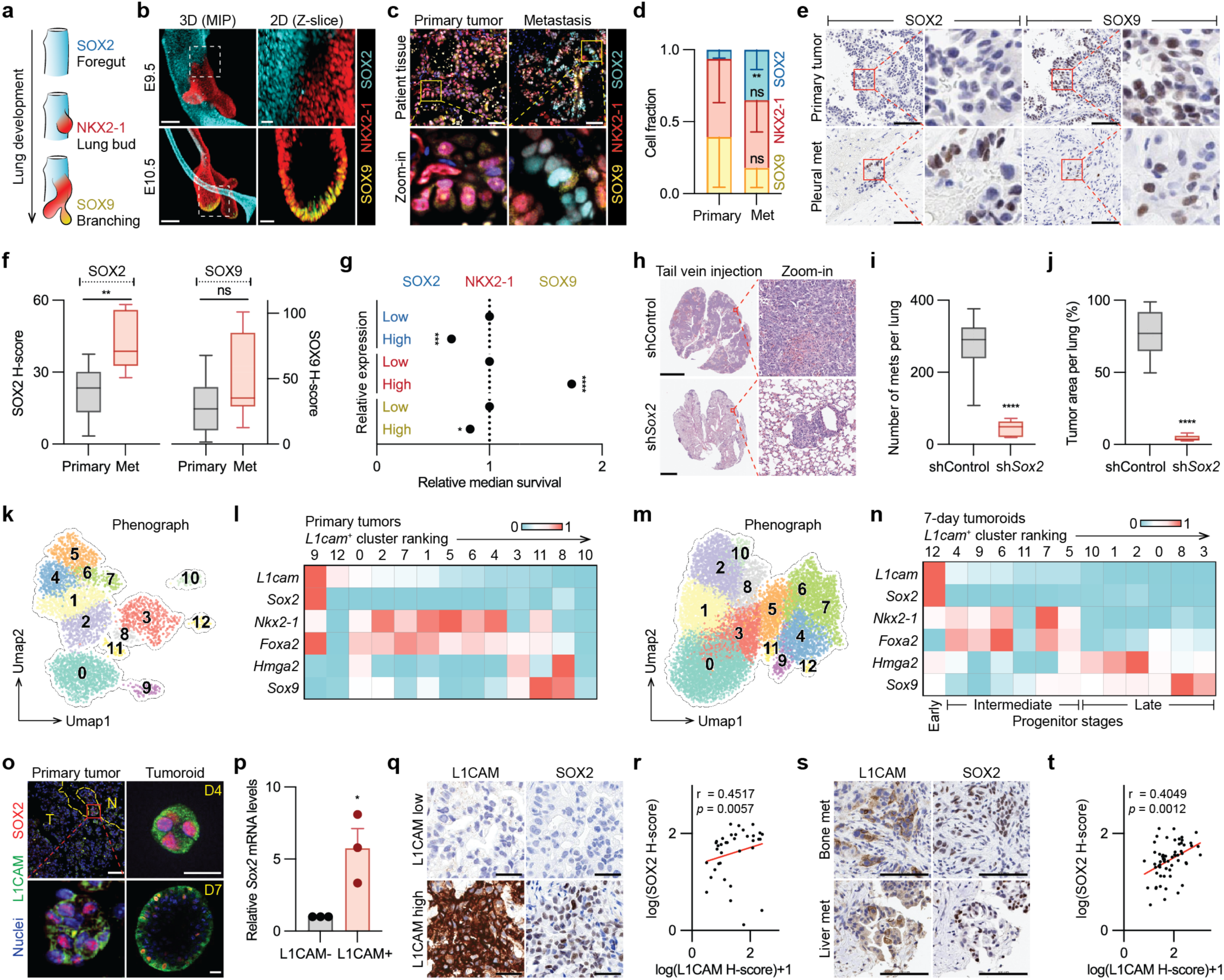
L1CAM marks LUAD SOX2^+^ progenitors. **a**, Schematic summary of lung developmental continuum from early to late progenitor stages defined by specific transcription factors. **b**, Representative 3D maximum intensity projections (MIP) images of SOX2, NKX2-1 and SOX9 IF in the anterior foregut domain in embryonic day E9.5 mouse embryos and the distal lung bud tip in E10.5 mouse embryos. The dashed box demarcates the SOX2/NKX2-1 boundary in E9.5 and NKX2-1/SOX9 boundary in E10.5 as illustrated in the zoomed 2D (Z-Slice) panels on the right. All data were independently validated from replicate samples of at least *n =* 4 embryos with similar results obtained. **c**, SOX2, NKX2-1, and SOX9 IF staining in patient-derived primary tumor and metastasis tissue sections. Magnified regions are highlighted in yellow boxes. Scale bar, 50 μm. **d**, Fraction of cancer cells expressing SOX2, NKX2-1, or SOX9 by IF analysis in patient-derived primary tumor and metastasis tissue sections. Mean ± S.D. *n* = 6 patient samples each. ns, *P* = 0.4848 (NKX2-1), 0.3095 (SOX9); \*\**P* = 0.0043. **e**, SOX2 and SOX9 IHC staining in LUAD patient-derived primary tumor and pleural metastasis samples. Magnified regions are highlighted in red boxes. Scale bar, 100 μm. **f**, Box and whisker plots of SOX2 and SOX9 IHC H-score in the patient-derived primary tumor and metastasis samples. Primary, *n* = 15; Metastasis, *n* = 7. ns, *P* = 0.1229; \*\**P* = 0.0011. **g**, Relative median survival of LUAD patients based on the expression of *SOX2*, *NKX2-1*, and *SOX9* in the primary tumor. Gene expression and survival data compiled from GEO, EGA and TCGA. *SOX2* (low, *n =* 706; high, *n* = 705); *NKX2-1* (low, *n =* 1083; high, *n* = 1083); *SOX9* (low, *n =* 1083; high, *n* = 1083). \**P* = 0.0219; \*\*\**P* = 0.0003; \*\*\*\**P* < 0.0001. **h**, H&E staining of lung tissue sections harboring metastatic colonies upon tail vein injection of KP tumoroid cells expressing control or *Sox2* short hairpins (sh). Magnified regions are indicated in red boxes. Scale bar, 5 mm. **i,j**, Number per lung (**I**) and percent area (**j**) of metastases in the experiment of panel (**h**). *n =* 10. \*\*\*\**P* < 0.0001. **k**, UMAP of scRNA-seq data from autochthonous KP tumors collected 32 weeks after viral instillation. *n* = 4,489 cells. Transcriptionally distinct clusters were computed by Leiden algorithm and numbered. **l**, Heatmap showing imputed average LUAD developmental marker expression in the clusters from primary tumors ranked left to right according to the average *L1cam* expression level. **m**, UMAP of scRNA-seq data from KP tumoroids collected after 7 days. *n* = 12,962 cells. Transcriptionally distinct clusters were computed by Leiden algorithm and are represented by a different color and a number. **n**, Heatmap showing imputed average LUAD developmental marker expression in the clusters from 7-day tumoroids ranked left to right according to the average *L1cam* expression level. Lung developmental progenitor stages represented by the predominant expression of *Sox2* (*early* developmental stage), *Nkx2-1* and *Foxa2* (*middle* progenitor stage), *Hmga2* and *Sox9* (*late* progenitor stage) are indicated. **o**, Representative L1CAM and SOX2 IF staining of the invasive front (*dotted yellow line*) within an autochthonous KP tumor (32 week post-Cre) and of KP-derived tumoroids grown for 4 days or 7 days. *T*, tumor; *N*, normal tissue. Magnified regions are indicated in red boxes. Scale bar, 50 μm for primary tumor; 20 μm for tumoroids. **p**, Relative *Sox2* mRNA expression of KP tumoroid-derived cells sorted by L1CAM expression. Mean ± S.E.M. *n* = 3. \**P* = 0.0258. **q**, L1CAM and SOX2 IHC staining of representative LUAD PDXs with high or low expression of L1CAM. Scale bar, 50 μm. **r**, Scatter plot and linear regression (*red line*) of L1CAM versus SOX2 IHC H-score data from LUAD PDXs. Data are log-transformed for visualization. *n =* 36. **s**, L1CAM and SOX2 IHC staining in serial sections of patient-derived bone metastasis and liver metastasis tissues. Scale bar, 100 μm. **t**, Scatter plot and linear regression (*red line*) of L1CAM versus SOX2 IHC H-score data from LUAD patient primary and metastasis samples. Data are log-transformed for visualization. *n =* 61 samples (primary, 15; metastasis, 46). Spearman correlation was used to calculate the relationship between L1CAM expression and SOX2 expression (**r,t**). Data are shown as a box (median ± 25-75%) and whisker (maximum to minimum values) plot (**f**,**i,j**). Statistical significance was assessed using the two-tailed Mann-Whitney test (**d**,**f**,**g**,**i**,**j**) or two-tailed *t* test after passing the Shapiro-Wilk normality test (**p**).

IF and IHC analysis of human LUAD samples showed that NKX2-1^+^ cells were abundant in low grade regions of primary tumor but were rare in high grade regions, whereas SOX2^+^ cells were enriched in high grade regions and in metastases (Fig. 2c-f and Extended Data Fig. 5a,b). In KP mice, SOX2^+^ cells were rare in primary tumors and enriched in autochthonous metastases, whereas SOX9^+^ were similarly abundant in primary and metastatic tumors (Extended Data Fig. 5c). Among the three lung developmental markers, patients with high *SOX2* expression split by the median expression value marked the worst clinical outcome in the integrated GEO, EGA, and TCGA databases (Fig. 2g)^41^.

To assess the role of SOX2, we used doxycycline-inducible expression of a sh*Sox2* hairpin in KP tumoroid cells to avoid global effects of a permanent SOX2 depletion (Extended Data Fig. 5d). These cells, and cells conditionally expressing a scrambled shRNA (shControl), were injected via the tail vein into immunodeficient NOD *scid* gamma (NSG) mice, which were fed doxycycline containing diet. The SOX2 knockdown markedly decreased the formation of pulmonary metastases (Fig. 2h-j), consistent with a role of SOX2^+^ cells as metastasis progenitors.

### SOX2^+^ LUAD cells show peak L1CAM expression

We conducted single-cell RNA sequencing (scRNA-seq) analysis of cancer cells freshly dissociated from KP primary tumors or grown as tumoroids over 7 days. Data analysis using the Leiden algorithm^42^ showed that cancer cells from primary tumors form 13 transcriptionally distinct clusters (Fig. 2k). *Sox2* was exclusively expressed in only one of these transcriptional cell clusters (Fig. 2l and Extended Data Fig. 5e). A ranking of the clusters based on *L1cam* expression levels showed peak expression of *L1cam* in the SOX2 cluster (Fig. 2l and Extended Data Fig. 5f,g). The SOX2 cluster and most of the other transcriptional clusters expressed a mix of developmental markers, in line with the observation that cancer cells in primary tumors from LUAD patients exhibit a mixed progenitor profile^6^.

KP tumor cancer cells grown as tumoroids for 7 days displayed 13 transcriptional cell clusters (Fig. 2m). *Sox2* was exclusively expressed in one cluster (cluster #12) and this cluster again showed peak expression of *L1cam* (Fig. 2n and Extended Data Fig. 5h-j). The 7-day organoid cells formed a more distinctive continuum of developmental stages than the primary tumor, comprising clusters that predominantly expressed *Sox2*, or *Nkx2-1* plus *Foxa2*, or *Hmga2,* or *Sox9* (Fig. 2n). This result mirrored the continuum of developmental stage markers observed in human LUAD metastasis^6^. Notably, ranking the 7-day organoid clusters based on their *L1cam* expression levels matched the developmental order of these stages, from the SOX2 cluster showing peak *L1cam* expression, the NKX2-1 and FOXA2 clusters with low *L1cam* expression, and the HMGA2 clusters and SOX9 clusters devoid of *L1cam expression* (Fig. 2n).

Confirming these results, IF analysis showed SOX2 staining in a subset of L1CAM^+^ cells within the micropapillary invasive front of KP primary tumors and within KP tumoroids (Fig. 2o). Sorted L1CAM^+^ KP tumor-derived cells expressed high levels of *Sox2* mRNA compared to L1CAM^−^ cells (Fig. 2p). A correlation between L1CAM and SOX2 expression was also evident in L1CAM^high^ versus L1CAM^low^ PDXs (Fig. 2q,r), and in L1CAM^+^ versus L1CAM^−^ areas of patient-derived primary tumors and metastases (Fig. 2s,t). Collectively, these results suggested that LUAD cancer cells that undergo natural or experimental dissociation and growth deploy a fetal lung developmental program including L1CAM^+^/SOX2^+^ primitive progenitors and cells expressing more developmentally advanced markers and lower or no L1CAM.

### L1CAM drives SOX2 expression in LUAD

The association between L1CAM and SOX2 expression in LUAD cancer cells was underscored by additional observations. Autochthonous KPL1 tumors lacked SOX2^+^ cells (Fig. 3a,b), and KPL1 tumoroids were devoid of *Sox2* mRNA expression, as shown by qRT-PCR analysis (Fig. 3c,d). When a high number (1 x 10^5^) of tumoroid cells was injected via the tail-vein of mice, KP tumoroids generated massive lung metastatic tumors containing SOX2^+^ and SOX9^+^ cancer cells whereas KPL1 tumoroids generated only small colonies containing SOX9^+^ cells but devoid of SOX2^+^ cells (Fig. 3e,f and Extended Data Fig. 5k).

**Figure 3.**
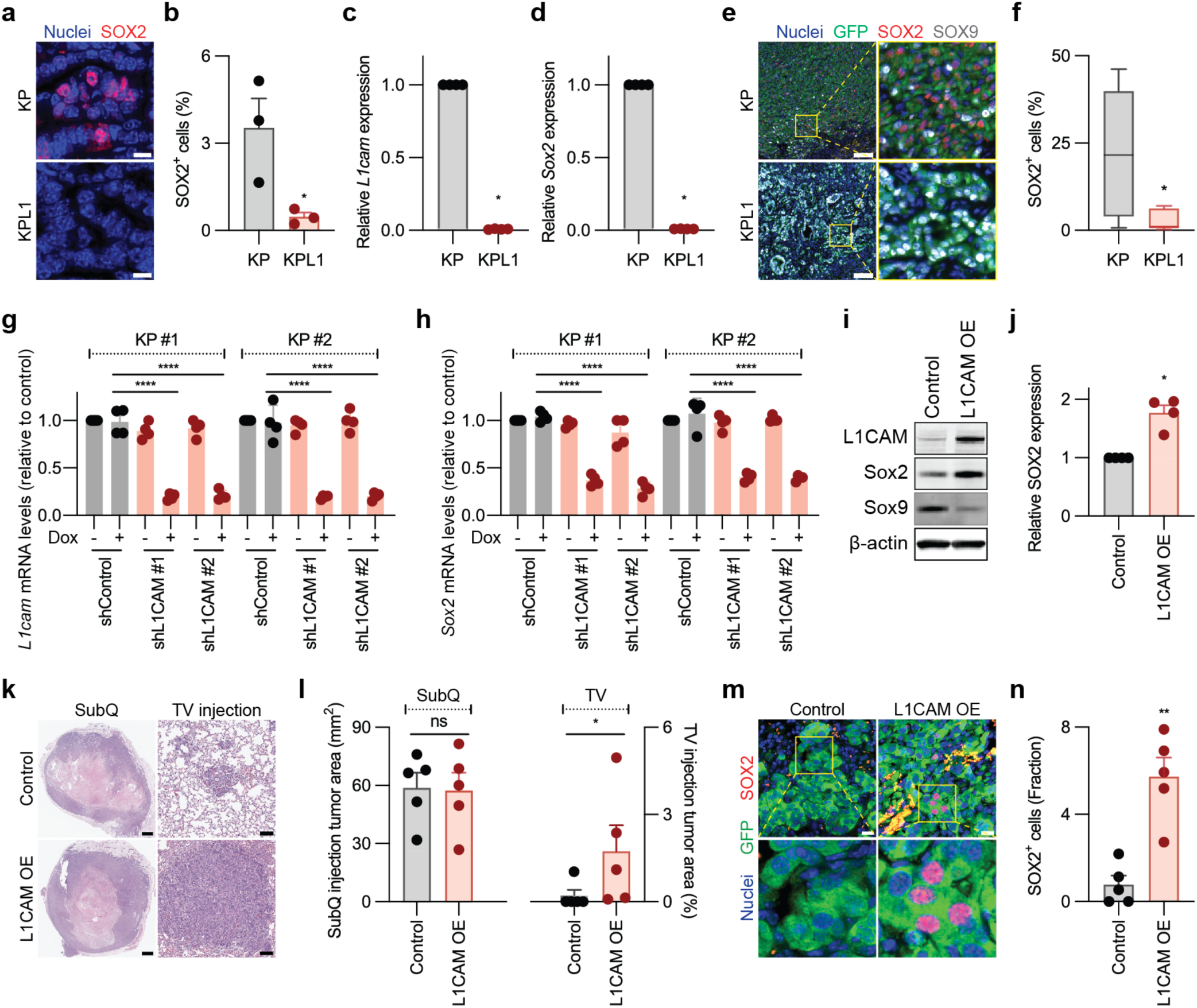
L1CAM promotes metastatic SOX2 expression. **a**, SOX2 IF staining of KP and KPL1 primary tumors (32 week post-Cre). Scale bar, 10 μm. **b**, Percentage of SOX2^+^ cells in the experiment from panel (**a**). *n* = 3 mice for each condition. \**P* = 0.0411. **c**,**d**, Relative *L1cam* mRNA expression and *Sox2* mRNA expression in KP and KPL1 tumoroids. KP, *n =* 4; KPL1, *n =* 4. \**P* = 0.0286 (**c**,**d**). **e**, IF images of KP and KPL1 tumors in the lung after injecting a high number (1 x 10^5^) of single cells from KP and KPL1 tumoroids into athymic mice via the tail vein. Magnified regions are highlighted in yellow boxes. Scale bar, 100 μm. **f**, Proportion of SOX2^+^ cells detected by IF in lung colonies after injecting a high number (1 x 10^5^) of KP or KPL1 cells via the tail vein in athymic mice. KP, *n =* 10; KPL1, *n =* 5. \**P* = 0.0400. **g**,**h**, Relative expression of *L1cam* (**g**) and *Sox2* (**h**) in two independent KP tumoroid lines upon Dox-dependent conditional knock down of *L1cam*. *n* = 4. \*\*\*\**P* < 0.0001. **i**,**j**, Western immunoblot analysis (**i**) and quantification (**j**) of SOX2 and SOX9 levels in control and L1CAM-overexpressing (OE) KP tumoroids. *n =* 4. \**P* = 0.0286. **k**,**l**, H&E staining (**k**) and quantification (**l**) of subcutaneous tumors or lung metastases after injection of KPL1 cells with or without L1CAM overexpression into athymic mice, analyzed at 4 weeks after subcutaneous (subQ) injection or 5 weeks after tail vein (TV) injection. Scale bar, 1 mm (SubQ) and 100 μm (TV). *n* = 5. ns, *P* > 0.9999; \**P* = 0.0238. **m**, GFP (cancer cells) and SOX2 IF staining of control vs L1CAM overexpressing KPL1 cells in lung metastasis after 5 week-post tail vein injection. Magnified regions are highlighted in yellow boxes. Scale bar, 10 μm. **n**, Fraction of SOX2^+^ cells in the experiment from panel (**m**). Control, *n* = 5; L1CAM overexpression, *n* = 5. \*\**P* = 0.0079. Data are shown as a box (median ± 25-75%) and whisker (maximum to minimum values) plot (**f**). Bar graphs, mean ± S.D. (**c**,**d**,**g**,**h**,**j**) or ± S.E.M. (**b**,**l**,**n**). Statistical significance was assessed using the two-tailed *t* test after passing the Shapiro-Wilk normality test (**b**), two-tailed Mann-Whitney test (**c**,**d**,**f**,**j**,**l**,**n**) or one-way analysis of variance followed by the Tukey test (**g**,**h**).

To determine whether SOX2 expression requires L1CAM, we engineered KP tumoroid cells with doxycycline-inducible expression of shRNAs targeting *L1cam*. The conditional knockdown of L1CAM in tumoroids caused a reduction in *Sox2* expression and a concurrent increase in *Sox9* expression (Fig. 3g,h and Extended Data Fig. 5l), suggesting that L1CAM stimulates *Sox2* expression and delays the transition of these cells to a SOX9^+^ state. Overexpression of L1CAM increased the SOX2 level and decreased the SOX9 level in KP tumoroids (Fig. 3i,j). Overexpression of L1CAM in KPL1 tumoroids (Extended Data Fig. 5m) did not increase their capacity to grow as subcutaneous tumors but rescued their capacity to colonize the lungs after intravenous injection (Fig. 3k,l). The resulting metastases contained SOX2^+^ cells in contrast to the micrometastases formed by control KLP1 tumoroids (Fig. 3m,n). These results implicated L1CAM as an inducer of SOX2 expression for the promotion of metastasis.

### L1CAM interaction with the planar cell polarity complex

To determine how L1CAM drives SOX2 expression in LUAD cells, we queried our single-cell transcriptional dataset for signaling pathway signatures that were differentially expressed in the L1CAM^+^/SOX2^+^ KP cells compared to other transcriptional cell clusters. Analysis of three independent databases –the Gene Ontology (GO)^43^, Reactome^44^, and PANTHER^45^ databases– identified planar cell polarity (PCP) and non-canonical WNT signaling as the most differentially enriched pathway signatures in L1CAM^+^/SOX2^+^ cells from primary tumors and KP tumoroids (Fig. 4a,b and Extended Data Fig. 6a).

**Figure 4.**
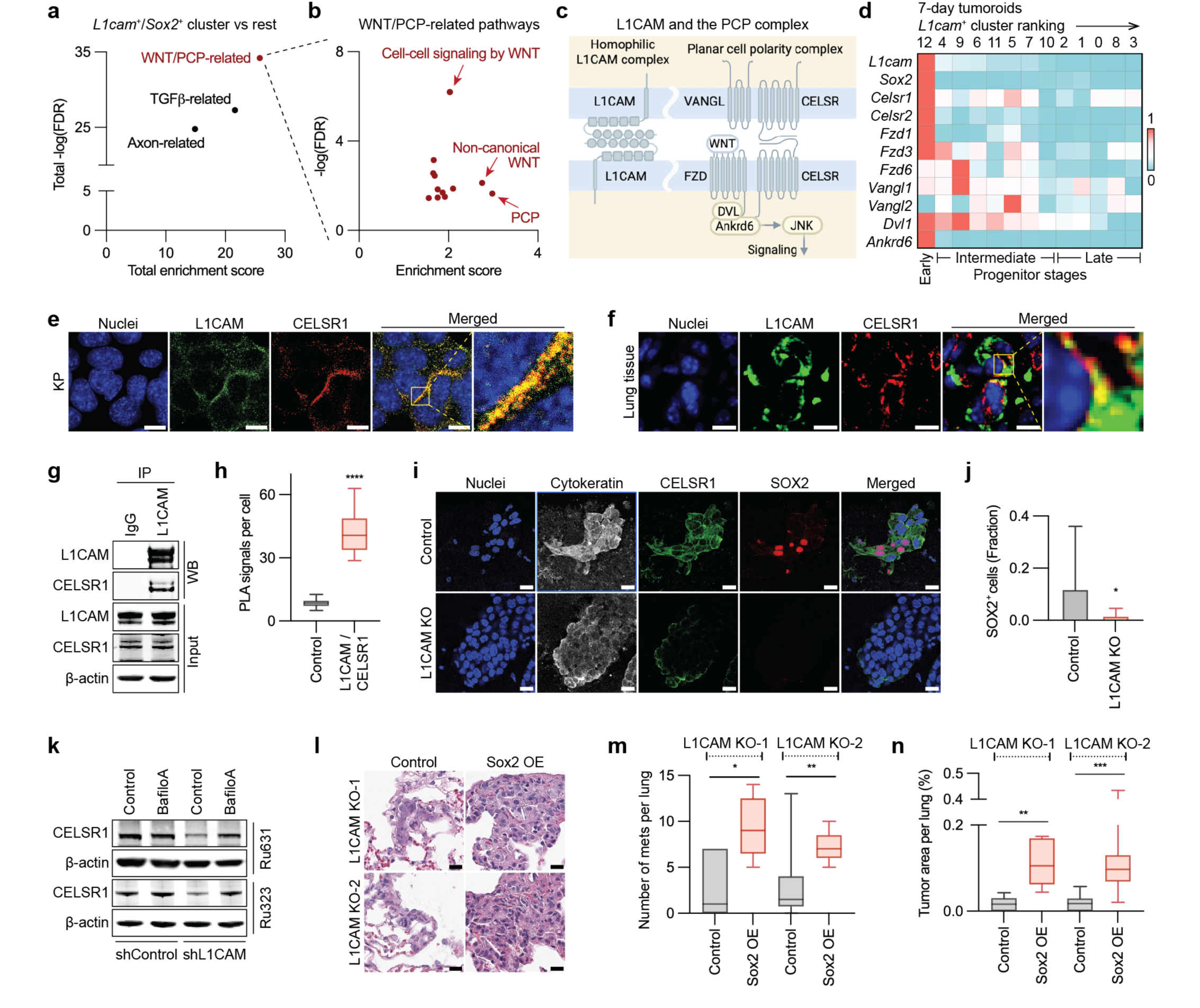
L1CAM enforces a SOX2^+^ metastasis-initiating state through PCP signaling. **a**, Scatter plot of accumulated enrichment score and negative log-transformed false discovery rate (FDR) of pathways associated with the L1CAM^+^/SOX2^+^ tumoroid cell cluster defined by scRNA-seq. The pathway with the highest total enrichment score and the lowest FDR is labeled in red. **b**, Scatter plot showing the expanded list of individual WNT/PCP-related terms. **c**, Schematic representations of L1CAM and PCP complex components at a cell-cell junction. **d**, Heatmap displaying the average expression of core PCP components in the KP tumoroid cell clusters defined in Figure 2N. The clusters are ranked left to right according to the average *L1cam* expression level. **e**, L1CAM and CELSR1 IF staining in KP tumoroid-derived cell monolayer. The merged image is magnified for visualization (*red box*). Scale bar, 10 μm. **f**, L1CAM and CELSR1 IF staining of lung metastasis one week after tail vein injection of H23 LUAD cells into athymic mice. Scale bar, 10 μm. **g**, Co-immunoprecipitation of L1CAM and the PCP component CELSR1 in H23 LUAD cell lysates. **h**, Quantification of L1CAM/CELSR1 PLA dots per cell in H23 LUAD cells. Negative PLA control, *n =* 25 cells; L1CAM/CELSR1 PLA, *n =* 31 cells. \*\*\*\**P* < 0.0001. **i**, Control and L1CAM-knockout (KO) H23 cell monolayers were subjected to cytokeratin, CELSR1, and SOX2 IF staining. Scale bar, 20 μm. **j**, Fraction of SOX2^+^ cells quantified in control versus L1CAM-knockout H23 cell monolayers. Mean ± S.D. Control, *n =* 43 cells; L1CAM KO, *n =* 58 cells. \**P* = 0.0488. **k**, Western immunoblotting analysis of control and L1CAM-knockdown (*shL1CAM*) PDXs after incubating the cells with 1 μM bafilomycin A for 24 h. **l**, H&E staining of lung sections of athymic mice after tail-vein inoculation of L1CAM-knockout H23 cells with or without SOX2 overexpression. Scale bar, 20 μm. **m**,**n**, Box and whisker plots showing the number (**m**) and percent area (**n**) of metastatic lesions per lung in the experiments of panel (**l**). *n =* 5 for L1CAM KO-1; 10 for L1CAM KO-2. \**P* = 0.0317; \*\**P* = 0.0012 in (**m**). \*\**P* = 0.0079; \*\*\**P* = 0.0002 (**n**). Data are shown as a box (median ± 25-75%) and whisker (maximum to minimum values) plot (**h**,**m**,**n**). Statistical significance was assessed using the two-tailed Mann-Whitney test (**h**,**j**,**m**,**n**).

The PCP complex is an asymmetrical assembly of membrane proteins located at the interface of adjacent epithelial cells^28–30^. Core PCP components include CELSR (cadherin EGF LAG seven-pass G-type receptor, 1 or 2), which is present on both interacting cell membranes, the WNT receptor Frizzled (FZD1-3 or FZD6) on one cell membrane, and the tetraspanin-like protein VANGL (Vang Gogh-like) located in the apposed cell membrane^29, 46^ (schematically summarized, together with L1CAM, in Fig. 4c). Formation of the PCP complex triggers non-canonical WNT signaling through the FZD cytosolic partners disheveled (DVL1-3) and ankyrin repeat domain 6 protein (ANKRD6)^46, 47^. This induces JNK to phosphorylate and activate c-Jun, an AP-1 family transcription factor^47, 48^. Indeed, the core PCP components CELSR1 and 2, FZD1, 3 and 6, VANGL1, DVL1 and ANKRD6 were highly expressed in the L1CAM^+^/SOX2^+^ KP tumoroid cells compared to other tumoroid transcriptional clusters (Fig. 4d). A similar expression pattern was observed in the L1CAM^+^/SOX2^+^ cells directly isolated from KP primary tumors (Extended Data Fig. 6b)

We conducted protein colocalization studies to explore this further. CELSR1 was colocalized with L1CAM at cell-cell interfaces membranes in KP tumoroids and cell monolayers, as determined by IF analysis (Fig. 4e and Extended Data Fig. 6c). Image analysis using steerable filters to extract curvilinear image features provided further evidence of L1CAM and CELSR1 colocalization at cell-cell interfaces in KP cell monolayers (Extended Data Fig. 6d,e). To analyze the co-localization of L1CAM and CELSR1 in metastatic tumors, we used the KRAS- and TP53-mutant human LUAD cell line H23, which expresses L1CAM in about 90% of the cells^49^. Tail-vein injections of H23 cells into athymic mice resulted in lung metastatic colonies that consistently showed colocalization of L1CAM and CELSR1 (Fig. 4f). Immunoprecipitation experiments (Fig. 4g) and proximity ligation assays (PLA) (Fig. 4h and Extended Data Fig. 6f) demonstrated a physical interaction between L1CAM and CELSR1 in H23 cells.

The PCP complex is assembled by subcellular enrichment of its components, and is negatively regulated by endocytosis and lysosomal degradation of its unassembled components^30^. CRISPR-Cas9 knockout of L1CAM reduced the levels of CELSR1 and SOX2 in H23 cells (Fig. 4i,j and Extended Data Fig. 6g), and in the Ru631 and Ru323 PDX models (Fig. 4k). The expression of CELSR1 in these PDX models was rescued by incubation with the pharmacological inhibitor of lysosomal degradation, bafilomycin A (Fig. 4k), suggesting that the loss of CELSR1 upon depletion of L1CAM involves endocytosis and degradation of the protein. Overexpression of SOX2 rescued the metastatic potential of L1CAM knockout H23 cells in tail-vein injection assays (Fig. 4l-n and Extended Data Fig. 6h).

### L1CAM promotes SOX2 expression through planar cell polarity signaling

These results raised the possibility that L1CAM promotes SOX2 expression and metastasis through PCP signaling. Therefore, we investigated the interdependence of L1CAM, CELSR1, and FZD6 as regulators of SOX2 expression. In KP tumoroids, L1CAM knockdown diminished the level of FZD6 at cell interfaces (Fig. 5a,b), suggesting that L1CAM stabilizes FZD6 as it did with CELSR1. A knockdown of FZD6 inhibited SOX2 expression (Fig. 5c,d), as did L1CAM depletion (refer to Fig. 3e,f). In the human cell line HEK293T, which expresses CELSR1 and FZD6 but not L1CAM, IF staining for CELSR1 and FZD6 showed a diffuse distribution of both proteins. The enforced expression of L1CAM in these cells resulted in the accumulation and colocalization of CELSR1 and FZD6 with L1CAM at cell interfaces (Extended Data Fig. 6i-l). The results further implicated L1CAM in the engagement of these PCP components as mediators of SOX2 expression in LUAD cells.

**Figure 5.**
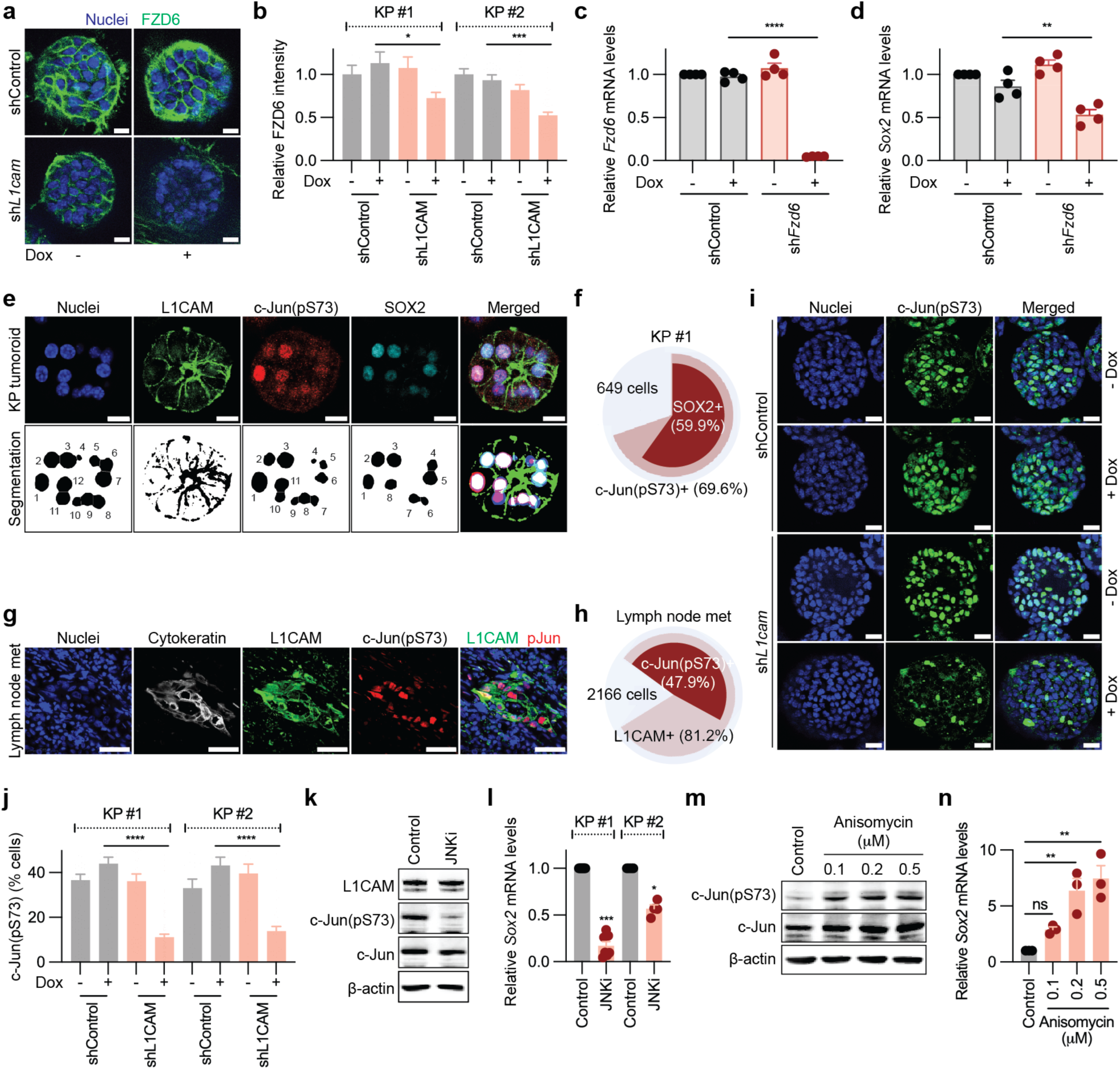
c-Jun drive SOX2 expression in L1CAM^+^ LUAD cells. **a**, FZD6 IF staining with counterstained nuclei in KP tumoroids upon conditional knockdown of *L1cam*. Scale bar, 10 μm. **b**, Relative intensity of FZD6 IF quantified in KP tumoroids upon conditional knockdown of *L1cam*. Left to right, *n =* 25, 23, 35, 29, 39, 41, 45, 53 tumoroids. \**P* = 0.0440, \*\*\**P* = 0.0010. **c**, Relative *Fzd6* mRNA level in upon conditional knockdown of *Fzd6* in KP tumoroids. *n =* 4 experiments. \*\*\*\**P* < 0.0001. **d**, Relative *Sox2* mRNA level upon conditional knockdown of FZD6 in KP tumoroids. *n =* 4 experiments. \*\**P* = 0.0040. **e**, IF staining for L1CAM, c-Jun(pS73), and SOX2 in KP tumoroids (*upper panels*) and image segmentation to quantify the signal (*bottom panels*). Scale bar, 10 μm. **f**, Pie chart showing the percent of cells staining positive for c-Jun(pS73) and SOX2 in KP tumoroids. *n =* 649 cells. **g**, IF staining for cytokeratin, L1CAM, and c-Jun(pS73) in a patient-derived LUAD lymph node metastasis. Scale bar, 50 μm. **h**, Pie chart showing the percent of cells staining positive for c-Jun(pS73) and L1CAM in patient-derived LUAD lymph node metastases. *n =* 2,166 cells from 12 different lymph nodes. **i**, c-Jun(pS73) IF staining and counterstained nuclei in KP tumoroids upon conditional knockdown of L1CAM. Scale bar, 20 μm. **j**, Percentage of cells staining positive for c-Jun(pS73) after conditional knockdown of *L1cam* in KP tumoroids. Left to right, *n =* 94, 67, 87, 101, 38, 41, 36, 36 tumoroids. \*\*\*\**P* < 0.0001. **k**, Western immunoblotting analysis of L1CAM, S73 phosphorylated and total c-Jun levels in H23 LUAD cells treated with 10 μM (JNK-IN-8) JNK inhibitor (JNKi) for 2 h. **l**, *Sox2* mRNA relative expression level upon incubation of KP tumoroids with 20 μM JNK inhibitor for 24 h. *n =* 7 for KP tumoroid #1; 4 for KP tumoroid #2. \*\*\**P* < 0.0006; \**P* = 0.0286. **m**, Western immunoblotting analysis of S73 phosphorylated and total c-Jun levels upon the treatments with anisomycin for 6 h. **n**, *Sox2* mRNA relative expression level upon incubation of KP cells with anisomycin for 6 h. *n =* 3. \*\**P* = 0.0064 (left), 0.0021 (right). The bar graph indicates mean ± S.E.M. (**b**-**d**,**j**,**l**,**n**). Statistical significance was assessed using the one-way analysis of variance followed by the Tukey test (**b**-**d**,**j**,**n**) or two-tailed Mann-Whitney test (**l**).

PCP/WNT signaling induces JNK-mediated phosphorylation c-Jun at serine 73^47, 48^. JNK-phosphorylated c-Jun (c-Jun(pS73)) was enriched in L1CAM^+^/SOX2^+^ KP tumoroid cells, as determined by IF (Fig. 5e,f). L1CAM and c-Jun(pS73) co-staining was also observed in LUAD patient-derived lymph node samples harboring incipient metastases (Fig. 5g,h). Knockdown of L1CAM decreased the levels of c-Jun(pS73) in KP tumoroid cells (Fig. 5i,j). Incubation of KP tumoroids with the JNK pharmacological inhibition JNK-IN-8^50^, decreased the level of c-Jun phosphorylation at pS73 and the expression of SOX2 without altering the levels of L1CAM or c-Jun (Fig. 5k,l). Conversely, pharmacological activation of JNK by incubation with anisomycin^51^ increased the levels of c-Jun(pS73) and the expression of *Sox2* (Fig. 5m,n). Collectively, these data indicated that L1CAM promotes the assembly and stabilization of the PCP complex at cell-cell interfaces, resulting in JNK-mediated activation of c-Jun as a mediator of *Sox2* expression.

### c-Jun and CHD1 jointly drive SOX2 expression in L1CAM^+^ LUAD cells

c-Jun is activated by various cues to regulate different aspects of cell behavior^52^. However, our results pointed at a specific role of c-Jun in regulating SOX2 expression in LUAD progenitors. These observations suggested the involvement of an additional factor(s) in the activation of SOX2 expression by L1CAM and PCP signaling. To test this hypothesis, we queried the differentially expressed genes from the L1CAM^+^/SOX2^+^ scRNA-seq cluster using the BART (binding analysis for regulation of transcription) algorithm^53^. This tool predicts functional transcriptional regulators based on the cis-regulatory regions of expressed genes. The 50 top ranked transcription factors were filtered based on chromatin immunoprecipitation (ChIP) enrichment analysis^54^ and ENCODE (encyclopedia of DNA elements) algorithms^55^. This integrated approach nominated CHD1 (chromodomain helicase DNA-binding protein 1) and MYC as factors of interest (Fig. 6a). MYC is a pleiotropic transcription factor that functionally interacts and shares target genes with SOX2 in the ChIP enrichment analysis database^54, 56^. Thus, the enrichment of a MYC transcriptional signature in L1CAM^+^/SOX2^+^ cells could reflect a functional overlap of MYC with SOX2 rather than a role of MYC in driving SOX2 expression.

**Figure 6.**
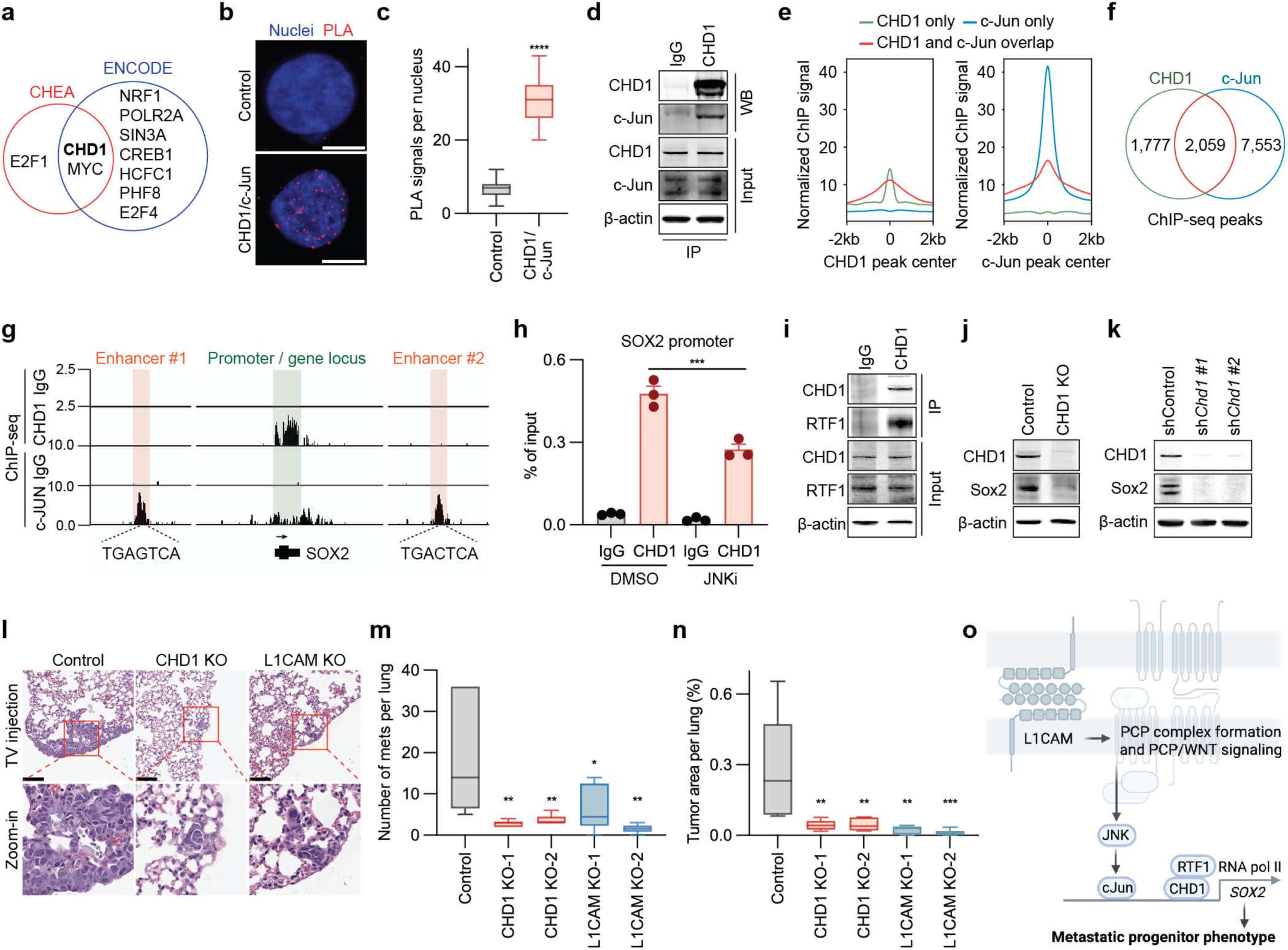
c-Jun and CHD1 jointly drive SOX2 expression in L1CAM^+^ LUAD cells. **a**, Venn diagram of transcription factors and chromatin modifiers differentially active in L1CAM^+^/SOX2^+^ KP tumoroid cells (scRNA-seq cluster #12, Figure 2N) based on CHEA and ENCODE databases. **b**, PLA image of CHD1 and c-Jun in H23 LUAD cells. The PLA signals are shown as red dots overlayed with nuclei staining. Scale bar, 10 μm. **c**, Quantification of PLA signals per nucleus. Control, *n =* 63 cells; CHD1/c-Jun, *n =* 39 cells. \*\*\*\**P* < 0.0001. **d**, Co-immunoprecipitation of CHD1 and c-Jun in KP tumoroids. **e**, Metaplots of CHD1 only (green), c-Jun only (blue), and CHD1/c-Jun overlap (red) ChIP-seq peak summits relative to peak center of CHD1 (left) or c-Jun (right) in H23 cells. **f**, Venn diagram showing the overlap between CHD1 and c-Jun genome-wide peaks. **g**, ChIP-seq analysis of CHD1 and c-Jun binding to the *Sox2* locus in H23 cells. *Sox2* enhancers (*red*), promoter and gene body (*green*) are indicated. **h**, ChIP-qPCR analysis of CHD1 binding to the *SOX2* promoter in H23 cells with and without JNK inhibitor. *n =* 3 experiments. \*\*\**P* = 0.0002. **i**, Co-immunoprecipitation of CHD1 and the transcriptional elongation factor RTF1 in H23 LUAD cells. **j**, Western immunoblotting analysis of SOX2 levels upon CHD1 knockout in H23 LUAD cells. **k**, Western immunoblotting analysis of SOX2 levels upon CHD1 knockdown in KP tumoroids using two different shRNAs. **l**, H&E staining of lung sections from NSG mice after tail-vein inoculation of H23 cells with CRISPR/Cas9-induced knockouts of CHD1 or L1CAM. Magnified regions are shown (*red boxes*). Scale bar, 100 μm. **m**,**n**, Box and whisker plots showing the number (**m**) and percent area (**n**) of metastatic lesions per lung in the experiments of panel (**j**). *n =* 6 per experimental condition. \*\**P* = 0.0035, 0.0060, 0.0015 from left to right; \**P* = 0.0301 in (**m**). \*\**P* = 0.0026, 0.0028, 0.0011 from left to right; \*\*\**P* = 0.0006 in (**n**). **o**, Model of L1CAM-dependent PCP activation of c-Jun/CHD1 driven SOX2 expression in LUAD progenitors to generate a metastasis-initiating state. Data are shown as a box (median ± 25-75%) and whisker (maximum to minimum values) plot (**c**,**m**,**n**). The bar graph indicates mean ± S.E.M. (**h**). Statistical significance was assessed using a one-way analysis of variance followed by the Tukey test (**h**,**m**,**n**) or two-tailed Mann-Whitney test (**c**).

Interestingly, CHD1 is a DNA-binding ATPase and chromatin remodeling factor^57^ and promotes pluripotency in stem cells^58^. PLA and co-immunoprecipitation experiments in H23 cells demonstrated an interaction between CHD1 and c-Jun in the nucleus (Fig. 6b-d). ChIP and sequencing (ChIP-seq) analysis revealed that CHD1 and c-Jun bind to common genomic regions in H23 cells (Fig. 6e,f). *SOX2* showed binding of c-Jun to enhancer sites containing the AP-1 binding motif [5’-TGA(G/C)TCA-3’] as well as broad binding across the *SOX2* promoter region and gene body (Fig. 6g). CHD1 ChIP-seq analysis showed specific binding to the *SOX2* promoter and gene body (Fig. 6g). Cell incubation with JNK inhibitor reduced CHD1 binding to the *SOX2* promoter, as determined by ChIP-PCR analysis (Fig. 6h).

In yeast, CHD1 facilitates active transcription by directly interacting with elongation factor RTF1, a component of the Paf1 complex that accompanies RNA polymerase II (RNAPII) during all stages of transcription^59^. Indeed, immunoprecipitation of CHD1 showed an interaction with RTF1 (Fig. 6i). This aligns with the observed binding of CHD1 across the *SOX2* gene body.

CRISPR/Cas9 knockout of CHD1 in human H23 cells and mouse KP tumoroids with different shRNAs caused a decrease in SOX2 expression (Fig. 6j,k) and impaired the metastatic activity (Fig. 6l-n), mirroring the effects of knocking down L1CAM, FZD6, or SOX2. However, SOX2 overexpression rescued poorly the metastatic activity of CHD1 knockout cells (Extended Data Fig. 6m,n), possibly due to the pleiotropic role of CHD1 as a chromatin remodeling factor.

In sum, these results collectively suggest that L1CAM-dependent activation of PCP/WNT signaling activates c-Jun to promote *SOX2* expression and a metastatic progenitor phenotype with CHD1 in LUAD cells (Fig. 6o).

## Discussion

In this study, we identify L1CAM as a critical regulator of the SOX2⁺ progenitor state in lung adenocarcinoma (LUAD), a cell state with the capacity to initiate multi-organ metastasis. We elucidate a mechanistic link between L1CAM and planar cell polarity (PCP) signaling through JNK activation, which drives SOX2 expression in these metastatic progenitor cells.

Primary LUAD tumors contain a heterogeneous population of cancer cells co-expressing markers such as NKX2-1, FOXA2, HMGA2, and SOX9, consistent with previous findings in human tumors^6^ and indicative of phenotypic plasticity^5–7^. While SOX2^+^ cells are rare in primary tumors, they are consistently enriched in metastatic lesions and co-exist with other progenitor cells expressing stage-specific markers that recapitulate fetal lung development. This developmental continuum is evident in tumoroids generated from KP mouse tumors, where early SOX2^+^ states are already established by day 7 in culture. Functionally, SOX2 is necessary for metastasis: its knockdown suppresses LUAD metastatic potential, while its overexpression rescues metastasis in L1CAM-deficient cells. These findings support a model in which LUAD tumors progressively generate a spectrum of progenitor stages, culminating in a SOX2^+^ metastatic progenitor with tumor-regenerative capabilities. This transition may occur either before or after dissemination. Supporting this idea, HMGA2^+^ cells, previously implicated in epithelial-mesenchymal transition and metastasis, may either generate earlier progenitor intermediates or serve distinct pro-metastatic functions^60, 61^.

Our data establish a link between L1CAM and SOX2 expression in LUAD cells. While L1CAM is absent from normal lung epithelium, it is expressed in LUAD tumors from both patients and KP mice and is primarily localized at the invasive front. Its expression, which is induced under conditions of adherens junction disruption^9^, is enriched in metastatic contexts, including patient-derived pleural fluids, distant metastases, PDXs, and autochthonous metastases in GEMMs. SOX2^+^ cells from both KP tumors and tumoroids exhibit high L1CAM expression, marking a distinct L1CAM^+^/SOX2^+^ state, which contrasts with the lower L1CAM expression in cells expressing intermediate (NKX2-1, FOXA2) or advanced (HMGA2, SOX9) progenitor markers. L1CAM is dispensable for the initiation of primary LUAD tumors. Tumors lacking L1CAM in KPL1 mice are indistinguishable from those in KP mice in terms of initial tumor burden, growth kinetics, histological grade, and capacity for subcutaneous growth. However, KPL1 mice lack spontaneous metastases, and KPL1 tumoroids fail to form metastases from the circulation, indicating that L1CAM is required for metastatic colonization. Moreover, L1CAM is required for SOX2 expression.

Mechanistically, we define a L1CAM–PCP–JNK–SOX2 signaling axis. L1CAM^+^/SOX2^+^ cells display upregulation of PCP components and activation of JNK. We show that L1CAM physically interacts with CELSR1, stabilizing it and FZD6 at cell-cell interfaces. This leads to JNK activation and phosphorylation of c-Jun, which in turn binds to a *SOX2* enhancer and cooperates with CHD1 and RTF1 to drive *SOX2* transcription. In the absence of L1CAM, CELSR1 becomes destabilized, JNK/c-Jun signaling is impaired, and SOX2 expression declines. This mechanism is distinct from our previously identified role of L1CAM in post-extravasation capillary cooption. In that context, L1CAM promotes the initial proliferation of extravasated metastatic cells by mediating binding to perivascular basement membrane and activating YAP and MRTF transcription factors via β1 integrins and integrin-linked kinase^27^. Thus, L1CAM plays a dual role as promoter of LUAD metastasis, a role in the sustained expression of SOX2 progenitor identity via PCP signaling and role in the initiation of proliferation of extravasated metastatic cells through integrin signaling. Both roles may coexist within the same metastatic lesion. This role of PCP/FZD-linked non-canonical WNT signaling is distinct from, but likely synergistic with that of canonical WNT signaling previously implicated in LUAD progression^62, 63^.

PCP signaling could have broader relevance in other malignancies where L1CAM expression correlates with poor clinical outcomes^64–72^. These findings underscore the therapeutic potential of targeting L1CAM^+^ metastatic progenitors. Although L1CAM expression peaks in SOX2^+^ cells, it is also detectable in other LUAD progenitor subsets. Thus, L1CAM-directed therapies may circumvent the challenge of intratumoral heterogeneity by simultaneously targeting multiple progenitor populations.

## Acknowledgements

We thank the MSKCC Integrated Genomic Operation, Molecular Cytology Core Facility, Flow Cytometry Core Facility, Pathology Core Facility, Single Cell Analytics Innovation Lab, and Animal Imaging Core for their technical assistance. We especially thank FE Kombak and M Jain for IHC analysis, L Tian for lentivirus vector generation, and J. Lee for Ru631 PDX tumoroids expressing luciferase. This work was supported by National Institutes of Health grants R35-CA252978 (JM), P01-CA129243 (JM), R01-CA270116 (TT), R35-CA263816 (CMR), R01-DK084391, R01-HD094868 (AKH), P30-CA008748 (MSKCC), grants from the Alan and Sandra Gerry Metastasis and Tumor Ecosystems Center at MSK (JM), fellowships from the Josie Robertson Foundation (TT), the Rita Allen Foundation (TT), the Translational Research Oncology Training Program at MSKCC (JSP), and the Canadian Institute Health Research PDF Fellowship (MIG).

## Author contributions

JSP, YK and JM conceived the study and designed the experiments; JSP, YK, CK and LH performed molecular cell biology experiments, generated tumoroids, and performed metastasis assays; RS and RC performed scRNA-seq; MIG and AKH conducted whole embryo imaging analysis; XZ performed automated deep neural tumor grading; UKB provided histopathological grading of tumors; RPK performed epigenetics analysis; TT provided oversight in GEMM tumor generation and analysis; CMR provided PDXs and advice during the study; JSP, YK, and JM wrote the manuscript. All authors edited and approved the manuscript.

## Materials & Correspondence

Any additional information and related data are available from the corresponding authors upon reasonable request. No restriction on data availability applies.

## Tables of Supplemental content

- Supplement table 1: Sequences for sgRNA oligos used in CRISPR-Cas9 mediated knockout
- Supplement table 2: Sequences for SMARTvectors Mouse Inducible Lentiviral shRNA
- Supplement table 3: qPCR probes and sequences

## Methods

### Animal studies

All animal experiments were conducted in accordance with protocols approved by the MSKCC Institutional Animal Care and Use Committee (IACUC) and the Research Animal Resource Center (RARC) (IACUC Protocol #99-09-032). All mice were regularly monitored by the investigators and veterinary staff at the Research Animal Resource Center at MSKCC, with food and water provided ad libitum. Kras^LSL-G12D^ (B6.129S4-Kras^tm4Tyj^/J; Jackson Laboratory Cat# 008179, RRID: IMSR_JAX:008179), *Trp53*^fl/fl^ (B6.129P2-Trp53^tm1Brn^/J; Jackson Laboratory Cat# 008462, RRID: IMSR_JAX:008462), *Rosa26*^LSL-Cas9-EGFP^ (B6J.129(B6N)-Gt(ROSA)26Sor^tm1(CAG-cas9*,-EGFP)Fezh/^J; Jackson Laboratory Cat# 026175, RRID: IMSR_JAX:026175), NSG (NOD.Cg-Prkdc^scid^IL2rg^tm1Wjl^/SzJ, Jackson Laboratory Cat# 005557, RRID: IMSR_JAX:005557), C57BL/6J (Jackson Laboratory Cat# 000664, RRID: IMSR_JAX:000664), and B6(Cg)-Tyr^c-2J^/J (B6-albino, Jackson Laboratory Cat# 000058, RRID: IMSR_JAX:000058) mouse strains were obtained from The Jackson Laboratory. Athymic nude (*Foxn1^nu^*, Envigo Cat# 069, RRID: IMSR_ENV:HSD-069) mice were obtained from Envigo. *L1cam^fl/fl^*mice were provided by M. Schachner (Rutgers University). KP- and KPL1-Cas9 mice were generated on C57BL/6J background. The genotype of KP and KPL1 mice was validated by Transnetyx from clipped tails (1 mm). Animals were maintained at temperatures of 21.1–22.2 °C, 30–70% humidity, 10–15 fresh air exchanges hourly, and a 12:12 h light / dark cycle (lights were on from 06:00–18:00).

#### Lenti-Sftpc-ffLuc-P2A-Cre vector

The Lenti-*Sftpc*-ffLuc-P2A-Cre vector was generated from a 317 bp Sftpc promoter sequence that drives gene expression in AT2 cells^73, 74^, a *ffLuc* template (Addgene #19166) and a *Cre* template (Addgene #60224). All three components were amplified by PCR using Q5 High-Fidelity Master Mix (NEB, M0492S). The pLL3.7 plasmid (Addgene #11795) was linearized with NsiI and PciI, and the fragments were assembled using Gibson Assembly Master Mix (NEB, #E2611).

#### LUAD tumor initiation

LUAD was conditionally generated in KP and KPL1 mice as previously described^32^. Briefly, female mice (8-10 weeks old) were administrated 2.5 x 10^4^ transducing units (TU) of pLenti-*Sftpc*-*ffLuc*-P2A-*Cre* via intratracheal instillation. Tumor initiation was monitored monthly and metastatic growth was monitored weekly in mice anesthetizing with 2-3% isoflurane/O_2_ and subjected to bioluminescence imaging (BLI) after retro-orbital injection of D-luciferin (150 mg/kg). BLI was performed in an IVIS Spectrum Xenogen instrument (PerkinElmer) and data were analyzed using Living Image Software v.4.5 (PerkinElmer). When imaging C57BL/6J mice, depilatory creams were used to remove any hair covering the region of interest before imaging.

Primary LUAD tumors were harvested at 26-32 weeks post lentiviral transduction or as specified. To isolate primary tumor cells, mice were euthanized and perfused with PBS through the left ventricle of the heart. Micro-dissected tumor tissues using a sterile scalpel were further dissociated by a mixture of dispase II (0.6 U/mL; Sigma Aldrich Cat# D4693), collagenase type IV (0.166 U/mL; Thermo Fisher Scientific Cat# 17104019), and DNase I (10 U/mL; STEMCELL Technologies Cat# 07900) in Dulbecco’s Modified Eagle’s Medium (DMEM) containing gentamicin (2 μg/mL; Gibco Cat# 15710064) at 37°C for 30-60 min. The cell suspension was filtered through 70 μm EASYstrainer (Greiner Bio-one) and spun at 300 g for 5 min at 4^°^C. The supernatant was aspirated, and red blood cells were lysed using RBC Lysis Buffer (Invitrogen Cat# 00-4333-57). Cells were then washed with PBS and spun at 300 g for 5 min at 4^°^C, re-suspended in ice-cold Fluorescence-Activated Cell Sorting (FACS) buffer [2% fetal bovine serum (FBS) in phosphate-buffered saline solution (PBS)], and strained through a 35 μm filter. Sorted GFP^+^ LUAD cells were seeded in a mixture of 50% Matrigel (Corning, Cat# CB-40230C) and 50% LUAD media (Advanced DMEM/F12; Gibco Cat# 12634-010), 2% of FBS, 2mM GlutaMAX (Gibco Cat# 35050061), 10 mM HEPES, 2 μg/mL of gentamicin, and 100 units/mL of penicillin/streptomycin).

#### Micro-CT imaging

CT images of mouse lungs were performed on the INVEON MicroPET-CT (Siemens Healthcare GmbH.Erlangen, Germany). A small-animal scanner was used to include the lung field with enhanced soft tissue contrast. Mice were anesthetized with 2-3% isoflurane/O_2_ throughout the procedure. The specific parameters of the X-ray tube were set up at a voltage of 70 kV, a current at 500 µA, and exposure time of 275 ms for each frame rotation. The projection settings used a half-gantry rotation, resulting in an image taken with 360 projections that lasted approximately 404 seconds total, for each CT. A Houndsfield (HU) scaling factor was applied during the reconstruction in real time, as well as the Shepp-Logan filter and a beam-hardening correction.

#### In vivo metastasis assays and limiting dilution assays

Female C57BL/6J (Jackson Laboratory 000664, RRID: IMSR_JAX:000664), athymic nude mice (Envigo 069, RRID: IMSR_ENV:HSD-069), NSG (Jackson Laboratory Cat# 005557, RRID: IMSR_JAX:005557) at 4 to 8 weeks of age were used for *in vivo* xenograft experiments. For multi-organ metastasis assays, 5 x 10^4^ cells were resuspended in 200 μL of PBS and inoculated into the right cardiac ventricle of athymic mice with a 26-gauge needle syringe. *Ex vivo* BLI of lung and kidney was measured 3 weeks after intracardiac injection. For lung colonization assays, 2 x 10^4^ cells were resuspended in 100 to 200 μL of PBS and inoculated into the lateral tail vein of athymic or NSG mice using a 28-gauge needle syringe. *Ex vivo* BLI of lung was performed 5 weeks after tail-vein injection of test cells. For intratracheal delivery, 2 x 10^4^ cells were resuspended in 100 μL of S-MEM medium (Gibco Cat# 11380037). *Ex vivo* BLI of lung was performed 5 weeks after intratracheal injection of KP or KPL1 cells. To compare the seeding ability of KP and KPL1 cells, athymic mice were injected with either cell line 1 x 10^5^ cells via tail veins and lungs were collected for analysis after 1 week.

To determine the role of SOX2 in metastasis, NSG mice were inoculated with KP cells engineered to conditionally express shRNAs targeting *Sox2* (sh*Sox2*) or a scrambled shRNA as control (shControl) into the lateral tail vein, at 10^5^ cells per mouse. Mice were fed a doxycycline diet (2500 mg/kg; ENVIGO TD.07383) throughout the duration of experiments. Four weeks after injection, mice were euthanized, and the lungs were harvested and fixed in 4% paraformaldehyde (PFA) overnight for histology analysis.

For limiting dilution assays, athymic mice were injected with KP or KPL1 cells subcutaneously at various doses from 5 x 10^4^ to 50 cells in PBS with Matrigel (1:1 ratio). After 6 weeks the mice were observed to determine whether subcutaneous tumors had developed. For KPL1 with or without L1CAM overexpression, subcutaneous tumors were examined after 4 weeks.

#### Whole-embryo IF-iDISCO clearing and imaging

Mouse embryos were dissected from E8.75-E12.5 in PBS and fixed in 4% PFA overnight at 4°C. Following three 10 min washes in PBS, embryos were gradually dehydrated for 1h in MeOH: 20% MeOH/PBS, 40% MeOH/PBS, 60% MeOH/PBS, 80% MeOH/PBS, and 100% MeOH twice at room temperature. At this stage, embryos were stored in 100% MeOH long-term at -20°C until needed for staining. When ready, embryos were first treated with 5% H_2_O_2_-methanol overnight at 4°C followed by rehydration steps: 80% MeOH/PBS, 60% MeOH/PBS, 40% MeOH/PBS, 20% MeOH/PBS, and PBS. Next, embryos were permeabilized in PBS, 0.2% Triton-X, 2.3% glycine and 20% DMSO overnight at 37°C. Samples were then transferred into blocking solution containing PBS, 0.2% Triton-X, 6% donkey serum, 10% DMSO overnight at 37°C. Embryos were then incubated for 48h at 37°C with primary antibodies at 1:200 (anti-SOX2 rat antibodies [Invitrogen Cat# 14-9811-82]; anti-NKX2-1 rabbit antibodies [Seven Hills Bioreagents Cat# WRAB-1231]; anti-SOX9 goat antibodies [R&D Systems Cat# AF3075]) diluted in PBS, 0.2% Triton-X, 0.001% heparin (PTwH), 3% donkey serum, and 5% DMSO. Samples were washed several times with PTwH and then incubated in secondary antibodies at 1:200 dilution (donkey anti-goat IgG Alexa Fluor^TM^ 488 [Thermo Fisher Scientific Cat# A-11055], donkey anti-rabbit IgG Alexa Fluor^TM^ 568 [Thermo Fisher Scientific Cat# A10042], and donkey anti-rat IgG Alexa Fluor^TM^ 647 [Thermo Fisher Scientific Cat# A78947]) for 48h at 37°C. Samples were washed several times with PTwH and stored at 4°C short-term prior to clearing. Embryos were gradually dehydrated for 1h in MeOH: 20% MeOH/PBS, 40% MeOH/PBS, 60% MeOH/PBS, 80% MeOH/PBS, and 100% MeOH, and stored at room temperature overnight. Samples were then incubated for 3h in 33% MeOH: 66% DCM (dichloromethane) followed by two 15 min washes with 100% DCM. Lastly, embryos were incubated with dibenzyl ether overnight and imaged on either Nikon A1R laser scanning confocal system or MuVi-SPIM Luxendo light-sheet system.

All image files were converted into ims. format using Imaris file converter and imported into Imaris software (Bitplane, Version 10.1.1). For better data management, embryos were cropped to specific region of interest and post-processing such deconvolution was performed one each image. Next, 3D surface volumes were rendered using the surface creation tab using the automatic threshold for each channel. Colors, brightness and contrast were adjusted accordingly to highlight key features in each image and the snapshot tool was used to capture the images. Final images were made in Adobe Illustrator CC (Adobe, Inc, CA).

### Patient-derived xenografts

LUAD PDXs were generated from primary and metastatic tumor samples as described ^75^. Briefly, samples were subcutaneously injected in the flank of a 6–8-week-old female mouse (Jackson Laboratory Cat# 005557, RRID: IMSR_JAX:005557). Tumors were harvested when the volume reached 1500 mm^3^ based on: V = ρχ/6 x L x W^2^ (L, length; W, width). Maximal tumor size was not exceeded. Serial passage was done by disaggregating and finely mincing the tumor, vigorously triturating in cold PBS, and straining through a 60 μm filter. Aliquots were cryopreserved in RPMI medium containing 10% DMSO.

### Cell lines and tissue culture

HEK293T control cells and HEK293T overexpressing L1CAM were cultured in DMEM with 10% FBS and 100 units/mL penicillin/streptomycin. H23 cells were purchased from American Type Culture Collection (ATCC) and cultured in RPMI with 10% FBS and 100 units/mL of penicillin/streptomycin. KP and KPL1 cells were cultured in the defined LUAD media or Advanced DMEM/F12 (Gibco Cat# 12634-010), 2% of FBS, 2 mM GlutaMAX (Gibco Cat# 35050061), 10 mM HEPES, 2 μg/mL of gentamicin, and 100 units/mL of penicillin/streptomycin. LUAD PDXs were cultured in RPMI with 10% FBS and 100 units/mL of penicillin/streptomycin. All cell lines were cultured at 37°C in a humidified incubator with 5% CO_2_. Prior to use, all cells tested negative for mycoplasma using the MycoAlert-PlusTM kit (Lonza Cat# LT07-710). When passaging, cells were detached and suspended as single cells in TrypLE Express Enzyme (Gibco Cat# 12604013) for 5 min at 37°C and neutralized by adding an equal volume of culture media before re-seeding. For inhibiting JNK signaling and c-Jun phosphorylation, cells were treated with JNK-IN-8 (10 μM, Selleck Chemicals Cat# S4901) dissolved in DMSO for 2 h before processing the samples for ChIP-seq. For quantifying Sox2 mRNA expression, KP tumoroids were treated with SP600125 JNK inhibitor (20 μM; Selleck Chemicals Cat# S1460) or DMSO for 24 h. When performing doxycycline-inducible shRNA targeting L1CAM or FZD6, KP tumoroids were incubated with or without 200 ng/mL of doxycycline for 2 days before proceeding to the next experimental steps.

#### Matrigel and tissue slice cultures

KP/KPL1 LUAD cells and PDXs were used to generate tumoroids. Cells were either directly dissociated into single-cell suspensions from tissues or suspended as single cells from a frozen stock. Once centrifuged, cells were resuspended in culture media and mixed with chilled Matrigel (Corning Cat# 356231) at 1:1 ratio before placement in a 35-mm glass-bottom dish (MatTek Cat# P35G-1.5-14-C). The glass-bottom area of dishes was initially coated with undiluted Matrigel solution to prevent cell attachment to the glass surface. Cells were then seeded on top of this Matrigel coating at 5 x 10^4^ cells in cold, diluted Matrigel solution. The mixture was incubated at 37°C for 30 min to promote gelation, and additional culture medium was added gently to the side of the cell-containing Matrigel domes. Cells were grown for 5-7 days or as described to generate mature tumoroids. For harvesting, the tumoroid-harboring Matrigel domes were transferred to a 15-mL conical tube and incubated in the Cell Recovery Solution (Corning Cat# 354270) for at least 30 min at 4°C on a rotator to digest Matrigel. Tumoroids were further dissociated into single cells using TrypLE for 15 min at 37°C, then diluted in culture media, centrifuged, and resuspended in fresh medium. Samples were then strained through a 35 μm filter to remove cell clusters and counted before additional experiments.

For tumoroid formation assay, KP/KPL1 cells (5 x 10^3^ cells/well) were plated in Matrigel and media mixtures at a 1:1 ratio onto a coated 96-well plate in a total volume of 100 μL. After gelation of Matrigel at 37°C for 30 min, an additional 100 μL of culture medium was gently added above the cells/Matrigel domes. The number of tumoroids was counted at day 7 and normalized by the number of cells seeded per well. Calcein AM (final concentration, 2 μM) was added per well, incubated for 20 min, and the tumoroids were imaged using the EVO fluorescence microscopy to measure the tumoroid cross-section area.

### Viral transductions

Lenti-*Sftpc*-*ffLuc*-P2A-*Cre* (see *Animal studies*) virus particles were generated in HEK293T cells transfected with 9 μg Lenti-*Sftpc*-*ffLuc*-P2A-*Cre* plasmid, 8 μg pSPAX2 (Addgene #12260), 3 μg pMD2.G (Addgene #12259) using Lipofectamine 3000 diluted in Opti-MEM media (Thermo Fisher Scientific) overnight. HEK293T cells were then cultured in DMEM containing 10% FBS (vol/vol), 2 mM L-Glutamine (Thermo Fisher Scientific), 100 U/ml Penicillin/Streptomycin (Thermo Fisher Scientific) for 48 h. Lentivirus containing media were collected by spinning at 300 g for 5 min at 4°C and straining through 0.45 μm Millex-HV filter (EMD Millipore). Lentivirus particles were concentrated using Amicon Ultra-15 centrifugal filter unit (Sigma Aldrich) according to the manufacturer’s instructions.

Lentivirus suspensions were generated by transfecting 70-80% confluent HEK293T cells with lentiviral vectors using the lentiviral packaging kit (Origene Cat# TR30037) or the Trans-Lentiviral shRNA packaging kit (Horizon Discovery Cat# TLP5912) according to the manufacturer’s instruction. Alternatively, lentivirus particles were produced from HEK293T cells with psPAX2 and pMD2.G (Addgene plasmids 12260 &12259) using Lipofectamine 3000. Viral particles were collected, filtered through 0.45 μm sterile filters, and incubated with cells of interest for 24 h with 8 μg/mL of polybrene before recovering cells in complete media. Lentivirus particles were concentrated using Amicon Ultra-15 centrifugal filter unit according to the manufacturer’s instructions.

Both KP/KPL1 tumoroids and PDX-derived tumoroids were transduced using TransDux MAX (System Biosciences Cat# LV860A-1) based on the manufacturer’s instruction. Briefly, tumoroids were isolated from Matrigel and dissociated into single cells as described above. Single cells were resuspended in RPMI media mixed with MAX Enhancer and TransDux solutions with 8mM HEPES. After adding lentivirus media, cells were transferred into a 24-well plate and spin-infected at 1500 g for 2 h at 32°C. After spinoculation, cells were resuspended in RPMI and Matrigel 1:1 solution and seeded back into a Matrigel-coated 35-mm glass-bottomed dish as tumoroid culture.

### CRISPR-mediated knockouts, overexpression, and shRNA knockdowns

KN2.0 CRISPR, non-homology mediated, knockout (KO) kits were used to generate L1CAM KO cells (Origene Cat# KN411601) and CHD1 KO cells (Origene Cat# N421575) according to the manufacturer’s instruction ^76^. Sequences for sgRNA oligos used in CRISPR-Cas9 mediated knockout are listed in Supplementary Table 1. The constructs were transfected into cells using TurboFectin (Origene Cat# TF81001), and cells were passaged for 2 weeks before isolating and expanding GFP^+^ single-cell knockout clones. Gene knockouts were validated by western immunoblotting of cell lysates.

Prior to performing metastasis assays, KP and KPL1 cells were transduced with a lentiviral TK-GFP-luciferase construct. For the constitutive expression of mouse *L1cam* and human *SOX2*, the constructs were commercially purchased and prepared (Origene Cat# MR211864 and Cat# RC200757L4). Products were transformed into stable competent Escherischia coli cells (New England Biolabs Cat# C3040H) according to standard protocols and spread on kanamycin (Fisher Scientific Cat# BP906-5) selective agar plates. Colonies were expanded, plasmid DNA was extracted using standard protocols and ZymoPURE II Plasmid Maxiprep Kit (Zymo Research Cat# D4203).

For shRNA-mediated protein knockdown, SMARTvectors Mouse Inducible Lentiviral shRNA (Horizon Discovery) were used for *L1cam*, *Sox2*, and *Fzd6* (see Supplementary Table 2 for sequences), in which expression is driven by a mEF1a promoter with a conditional TurboRFP reporter. The SMARTvector Inducible Non-targeting control was used as a negative control, and the Trans-Lentiviral shRNA packaging kit (Cat# TLP5912) was used to generate lentivirus. For knocking down L1CAM in PDXs, L1CAM human shRNA plasmid kit (Origene Cat# TL303597) was used. For knocking down *Chd1* in KP cells, Chd1 mouse shRNA plasmid kit (Origene Cat# TL516702) was used. ShControl cells were generated in both cases by transducing the cells with scrambled shRNA control (Origene Cat# TR30021). Viral transductions were performed as stated above, and stable cell lines were generated by puromycin selection for 2 weeks.

### Immunoblotting and Immunoprecipitation

Protein lysates were prepared in ice-cold RIPA buffer (Sigma-Aldrich Cat# R0278) supplemented with Halt protease and phosphatase inhibitor cocktail (Thermo Fisher Scientific Cat# 78442). Cells were ruptured by sonication, and the lysates were centrifuged and pelleted at 4°C. The supernatant was collected and quantified using the BCA protein assay (Thermo Fisher Scientific Cat# 23228) analyzed on a microplate reader (BioTek Synergy-H1) at 562-nm absorbance with Gen5 software (BioTek). Samples were diluted in reducing Laemmli SDS sample buffer (Thermo Fisher Scientific, Cat# J60015.AC) and boiled for 5 min. Proteins were separated on NuPAGE Novex 4-12% Bis-Tris gels (Thermo Fisher Scientific Cat# NP0322BOX) using 1x MOPS SDS running buffer (Thermo Fisher Scientific Cat# NP0001) and transferred to nitrocellulose membranes. Once membranes were blocked with Intercept Tris-buffered saline (TBS) blocking buffer (LICOR Biosciences Cat# 927-60001) for 1 h at room temperature, they were probed with antibodies against c-Jun (Cell Signaling Technology Cat# 9165, RRID: AB_2130165), c-Jun(pS73) (Cell Signaling Technology Cat# 9164, RRID: AB_330892), CHD1 (Cell Signaling Technology Cat# 4351, RRID: AB_11179073), β-actin (Sigma-Aldrich Cat# A1978, RRID: AB_476692), L1CAM (BioLegend Cat# 826701, RRID: AB_2564904), Sox2 (Cell Signaling Technology Cat# 3579, RRID: AB_2195767), CELSR1 (Bethyl Cat# A304-432A, RRID: AB_2620626), and RTF1 (Cell Signaling Technology Cat# 14737, RRID: AB_2798594) in blocking buffer for overnight at 4°C. After washing three times with TBS with 0.1% Tween (TBST), the membranes were probed with secondary antibodies (LICOR Biosciences) in Intercept TBS blocking buffer for 1 h at room temperature. After another wash, bands were detected with the 680 and 800 channels of an Odyssey CLx imager (LICOR Biosciences). Protein bands were quantified by measuring peak areas using ImageJ. The peak area for each protein was normalized against the peak area of a loading control.

For immunoprecipitation, protein lysates were prepared in ice-cold cell lysis buffer (Cell Signaling Technology Cat# 9803) supplemented with Halt protease and phosphatase inhibitor cocktail (Thermo Fisher Scientific Cat# 78442). Cells were sonicated, centrifuged, and pelleted at 4 °C, and the supernatant was collected. The samples were pre-cleaned with magnetic Dynabeads Protein G (50% bead slurry, Thermo Fisher Scientific Cat# 10003D) for 2 h or overnight at 4 °C to reduce nonspecific binding. The beads were pelleted using a magnetic rack, and the supernatant was collected and proceeded to immunoprecipitation. The supernatant was incubated with primary antibodies against L1CAM (Invitrogen, Cat# 14-1719-82, RRID: AB_891383), c-Jun (Cell Signaling Technology Cat# 9165, RRID: AB_2130165) or CHD1 (Cell Signaling Technology Cat# 4351, RRID: AB_11179073) overnight at 4°C on a rotator. Afterward, Dynabeads were incubated with immunoprecipitation samples at 4°C on a rotator for 2 h. The beads were then washed with cell lysis buffer five times. The pellets were reconstituted in a reducing Laemmli SDS sample buffer and boiled for 5 min before performing SDS–PAGE and immunoblotting.

For pharmacological treatment and immunoblotting, cells were treated with various agents prior to protein lysate preparations. 3 x 10^5^ cells were seeded in a 6-well plate and treated one day later with either DMSO or Bafilomycin A1 (Cell Signaling Technology Cat# 54645) for 24 hours at a concentration of 1μM. For activation of JNK signaling, cells were treated with various concentrations of Anisomycin (Tocris Cat# 1290) [0.1, 0.2, and 0.5 μM] or DMSO for 6 hours prior to protein lysate collection.

### Immunohistochemistry, immunofluorescence and microscopy

Lung cancer primary tumor and metastatic samples were obtained from patients confirmed Stage I, II, III or IV lung adenocarcinoma. Patient samples were selected based on the availability of a sufficient quantity of formalin-fixed paraffin-embedded tissue from patients who had given preoperative consent to tissue utilization for research purposes. Human specimens were obtained and approved under Memorial Sloan Kettering Cancer Center Institutional Review Board biospecimen research protocol 15-101. All patients provided pre-procedure informed consent. For staining of 4 μm sections, mouse metastatic tissue samples were fixed in 4% paraformaldehyde for 24 h followed by paraffin embedding. For tumoroid sections, Matrigel samples were first transferred to HistoGel and fixed in formalin for 24 h before paraffin embedding. Unstained slide sections were deparaffinized using Histo-Clear (National Diagnostics Cat# HS-200) and rehydrated in a series of ethanol washes. Antigen retrieval was performed in the Tris-based antigen unmasking solution (Vector Laboratories Cat# H-3300) in a steamer for 30 min. Ultra-High-Def Mouse on Mouse (MOM) blocking reagent (StatLab Cat# P1-M-100-HRP) was used for L1CAM staining in mouse tissues according to the manufacturer’s instructions. Sections were blocked in 10% normal goat serum (Life Technologies Cat# 50062Z) for at least 1 h at room temperature, and incubated with primary antibodies diluted in blocking solution overnight at 4°C.

For immunohistochemistry (IHC) staining, tissue sections were processed and stained with anti-L1CAM antibody (BioLegend Cat# 826701, RRID: AB_2564904), anti-Sox2 antibody (Cell Signaling Technology Cat# 14962, RRID: AB_2798664) or anti-Sox9 antibody (Cell Signaling Technology Cat# 82630, AB_2665492), via standardized automated protocols using a Ventana Discovery ST instrument. ImmPRESS horseradish peroxidase (HRP) anti-mouse IgG and ImmPACT DAB Peroxidase (Vector Labs) were used for detection. The slides were counterstained with hematoxylin, dehydrated and capped with a coverslip. Additional hematoxylin and eosin (H&E) staining was performed by Histowiz, Inc (Brooklyn, NY, US).

For immunofluorescence staining, samples were fixed in 4% PFA for 10 min and permeabilized with 0.5% of Triton X-100 in PBS for another 10 min. After incubating with 10% normal goat serum (Life Technologies Cat# 50062Z) for 1 h at room temperature, the samples were incubated with primary antibodies overnight at 4°C in blocking solution with antibodies against mouse L1CAM (Miltenyi Biotec Cat# 130-115-812, AB_2727206), human L1CAM (Santa Cruz Biotechnology Cat# sc-53386, RRID: AB_628937), Sox2 (Invitrogen Cat# 14-9811-82, RRID: AB_891383), CELSR1 (Millipore Sigma Cat# ABT119, RRID: AB_11215810), c-Jun(pS73) (Cell Signaling Technology Cat# 9164, RRID: AB_330892), GFP (Aves Labs Cat# GFP-1010, RRID: AB_2307313), mouse FZD6 (R&D Systems Cat# AF1526, RRID: AB_354842), human FZD6 (Abcam Cat# AB150545, RRID: AB_3697520), Sox9 (Invitrogen Cat# 14-9765-82, RRID:AB_2573006), Cleaved Caspase-3 (Cell Signaling Technology Cat# 9661, RRID: AB_2341188), anti-TTF1/NKX2-1 (abcam Cat# ab76013, RRID: AB_1310784), or human cytokeratin (Dako Cat# M3515, RRID: AB_2132885). After washing in PBS three times, samples were incubated with fluorophore-conjugated secondary antibodies for 1 h at room temperature in the dark, followed by another round of PBS washes and staining of nuclei with Hoechst 33342 (Thermo Fisher Scientific Cat# H3570). Sections were washed with PBS three times and mounted using ProLong diamond antifade mountant (Life Technologies Cat# P36970) before performing microscopy.

The H&E and IHC stained sample sections were digitally scanned using the Aperio AT2 slide scanner (Leica Biosystems) under 20x objective magnification. Scanned images of tissue sections were annotated for tumor compartments by pathologists. The membraneous or nuclear algorithm of Aperio ImageScope v12.4.3.7005 (Leica biosystems) was used under supervision of a pathologist to quantify the staining intensity of L1CAM and Sox2: 0+ (no staining), 1+ (weak staining), 2+ (moderate staining) or 3+ (strong staining). An H-score was assigned based on the intensity score and the percentage of scored cells: H-score = 0 x (% of cells with 0+ intensity score) + 1 x (% of cells with 1+ intensity score) + 2 x (% of cells with 2+ intensity score) + 3 x (% of cells with 3+ intensity score).

For fluorescence microscopy, slides were analyzed on a STELLARIS 8 confocal microscope (Leica), equipped with an Apo 63x 1.4 NA objective. Images were recorded with a Leica HyD (Leica Hybrid Detectors) controlled by Leica Application Suite X 4.5.0.25531 software. For scanning whole stained tissue sections, slides were scanned on a Pannoramic Scanner (3DHistech, Budapest, Hungary) using a 20x/0.8NA objective. Scans were visualized using Slideviewer (Version 2.7, 3DHistech). Live tumoroids were imaged by EVOS M5000 (Invitrogen). Fixed and stained images were also imaged from an Imager.Z1 epifluorescence microscope (Zeiss) with Apotome.2 optical sectioning module, equipped with the digital camera C11440 (ORCA-Flash4.OLT, Hamamatsu) and controlled by Zeiss ZEN 2.3 microscopy software.

### Image analysis

ImageJ 1.53n was used to detect and quantify the nuclei from Hoechst or IF staining of pJun (S73) and SOX2. Raw nuclei staining was first smoothened by a Gaussian Blur filter and segmented using pixel intensity thresholding. Overlapping nuclei were separated by a watershed method, and the number of nuclei was counted after filtering out particles that were smaller than 4 μm^2^ in size. Similarly, several metastatic lesions after injecting single-cell suspension of shControl KP tumoroids was quantified by pixel intensity thresholding of H&E staining images and separating clumped lesions by a watershed method. QuPath v0.5.1 was used to detect and quantify tumors from lung tissue sections. Tumor regions with the lung were manually segmented under the guidance of a pathologist, counted, and normalized by the total area of the lung. Both manual and automatic cell detection were used to quantify cells with positive IF staining.

L1CAM and CELSR1 images were analyzed by applying a steerable filter pipeline as previously described ^77^. In brief, image features were enhanced by multiscale steerable filtering and center lines of curvilinear features were captured. The junction fragments were divided into high- and low-confidence, and the fragments were connected by graphical matching to yield junction positions as a chain of connected pixels. From the detected junctions, the length and pixel intensity parameters were determined using customized scripts written in MATLAB 2018a (Mathworks). Junctions were shown as binary images, and the line thickness was enhanced by dilation using ImageJ for visualization.

For analysis of PLA images, nuclear or whole-cell boundaries in c-Jun/CHD1 or L1CAM/CELSR1 experiments, respectively, were manually segmented using ImageJ. The PLA fluorescence signal channel was set to threshold to avoid false-positive background signals. The number of PLA signal loci per nucleus or cell was determined based on the number of local fluorescence maxima.

### qRT-PCR analysis

RNA was extracted from tumoroid pellets using the RNeasy Mini Kit (Qiagen Cat# 74106). Total RNA (200 ng) was used to generate cDNA using the Transcriptor First-Strand cDNA synthesis kit (Roche Cat# 04379012001) or Superscript IV VILO Master Mix (Thermo Fisher Scientific Cat# 11756500). Relative gene expression was determined by quantitative PCR on the ViiA 7 Real-Time PCR System (Life Technologies) or QuantStudio 7 Pro (Thermo Fisher Scientific) using TaqMan gene expression assay probes (Life Technologies) or SYBR Green assays (Life Technologies) (see Supplementary Table S3 for probes and sequences). Relative mRNA levels were calculated by the 2−ΔΔCt method and were normalized to the expression of murine *Gapdh*.

### ChIP sequencing assays and data analysis

ChIP was performed using the SimpleChIP Enzymatic Chromatin IP Kit (Cell Signaling Technology Cat# 9003) according to the manufacturer’s instruction. Briefly, 10^7^ cells from a 15 cm tissue culture dish were crosslinked with 1% formaldehyde for 10 min at room temperature. The reaction was then quenched by adding glycine solution and incubating for 5 min at room temperature. After washing the cells two times with ice-cold PBS, cells were collected in ice-cold PBS supplemented with protease inhibitors (Cell Signaling Technology Cat# 7012) and centrifuged. Supernatants were aspirated, and cell pellets were snap frozen in liquid nitrogen and stored at -80°C until proceeding to the next experimental steps. Cells were lysed using the manufacturer’s buffer A (Cell Signaling Technology Cat# 7006) and resuspended in buffer B (Cell Signaling Technology Cat# 7007). Chromatin was digested using micrococcal nuclease (Cell Signaling Technology Cat# 10011) at 37°C for 20 min and the reaction was quenched by adding ETDA. Nuclei were centrifuged, resuspended in 1X ChIP buffer (Cell Signaling Technology Cat# 7008), and sonicated. Digested chromatins were incubated with designated antibodies at 4°C overnight on a rotator and incubated with pre-washed ChIP-Grade Protein G Magnetic Beads (Cell Signaling Technology Cat# 9006) for 2h at 4°C on a rotator. The samples were washed three times with low-salt ChIP buffer and one time with high salt ChIP buffer. Chromatin complexes were eluted by 1X ChIP elution buffer (Cell Signaling Technology Cat# 10009) for 30 min at 65°C with gentle vortexing, and reverse-crosslinked with Proteinase K (Cell Signaling Technology Cat# 10012) for 2 h or overnight at 65°C. DNA was purified using spin columns. For library construction and sequencing, DNA was quantified by PicoGreen and the size was evaluated by Agilent BioAnalyzer. The KAPA HTP Library Preparation Kit (Kapa Biosystems, Cat# KK8234) was used to prepare Illumina sequencing libraries with 0.02-5 ng of input DNA and 8-14 cycles of PCR. Barcoded libraries were run on the NovaSeq 6000 in a PE100 run, using the NovaSeq 6000 S2 or S4 Reagent Kit (200 or 300 Cycles) (Illumina). Approximately 40 million paired reads were generated per sample.

ChIP-seq sequencing reads were trimmed and filtered for quality and adapter content using version 0.4.5 of TrimGalore (https://www.bioinformatics.babraham.ac.uk/projects/trim_galore), with a quality setting of 15 and running cutadapt (v1.15) and FastQC (v0.11.5). Reads were aligned to human assembly hg38 with version 2.3.4.1 of bowtie2 (http://bowtie-bio.sourceforge.net/bowtie2/index.shtml) and were deduplicated using MarkDuplicates in version 2.16.0 of Picard Tools. To ascertain regions of target enrichment, MACS2 (https://github.com/taoliu/MACS) was used with a p-value setting of 0.001 and scored against matched IgG as the control. The BEDTools suite v2.29.2 (http://bedtools.readthedocs.io) was used to create normalized read density profiles. A global peak atlas was created by first removing blacklisted regions (http://mitra.stanford.edu/kundaje/akundaje/release/blacklists/hg38-human/hg38.blacklist.bed.gz) then merging all peaks within 500 bp for ChIP and counting reads with featureCounts (v1.6.1). Each signal was sequencing depth normalized to ten million uniquely mapped fragments. Peak-gene associations were created by assigning peaks to genes using linear genomic distance to transcription start site. Peak intersections were calculated using bedtools and intersectBed with 1 bp overlap. Profile plots were created using deepTools v3.3.0 by running computeMatrix and plotHeatmap on normalized bigwigs with average signal sampled in 25 bp windows and flanking region defined by the surrounding 2 kb.

### ChIP-qPCR analysis

DNA samples were immunoprecipitated and purified following the ChIP-seq protocol. Samples were analyzed by qRT-PCR using SimpleChIP universal qPCR master mix (Cell Signaling Technology Cat# 88989), and results were quantified as a percent of the total input chromatin using the equation: percent input = 2% x 2^(Ct^ ^2%^ ^input^ ^sample−Ct^ ^IP^ ^sample)^, where Ct is the threshold cycle of the PCR reaction.

### Single-cell RNA sequencing

LUAD cells were freshly isolated from mice and sorted by their GFP marker as described above. Freshly sorted cells were either sequenced directly as the sample day 0 or cultured in Matrigel as tumoroids for 7 days prior to sequencing. Tumoroids were collected and dissociated into single cells. Cells were sorted in ice-cold PBS with 0.04% BSA by flow cytometry using DAPI as a negative viability marker. Single-cell RNA-seq was performed on 10x Genomics Chromium platform as previously described^9^. The scRNA-seq libraries were prepared based on the manufacturer’s protocol for 3’ end reading. The individual transcriptomes of encapsulated cells were barcoded during RT step and resulting cDNA purified with DynaBeads followed by amplification per manual guidelines. Next, PCR-amplified product was fragmented, A-tailed, purified with 1.2X SPRI beads, ligated to the sequencing adapters and indexed by PCR. The indexed DNA libraries were double-size purified (0.6–0.8X) with SPRI beads and sequenced on Illumina NovaSeq S4 platform (R1 – 26 cycles, i7 – 8 cycles, R2 – 70 cycles or higher). The raw sequencing data were then demultiplexed and aligned to the mouse genome using 10X CellRanger pipeline. This generated an output matrix of mRNA counts per cell for every gene. We used the filtered matrix for all downstream analysis.

All processing of the resulting data was performed using the Scanpy package ^78^. For quality control, cells with less than 200 genes detected and genes expressed by less than 3 cells were filtered out. Based on the library distribution, cells with less than 1,000, greater than 5% mitochondrial RNA content or less than 5% ribosomal RNA content were excluded from analysis. Housekeeping genes MALAT1 and GM42418 were also removed, with 9,949 cells from freshly isolated KP tumor and 11,064 cells for day-7 KP tumoroids passing the filters.

For both sets of analysis (freshly isolated KP tumor and day-7 KP tumoroid samples), the data were normalized by library size followed by a log transformation (base = 2, pseudo count = 1). From the normalized count matrices, principal component analysis (PCA) was performed using the top 4,000 highly variable genes (HVGs). The selected principal components (PCs) were used to construct *k*-nearest neighbor graph (n_neignbors = 30) to generate uniform manifold approximation and projection (UMAP) layouts and to cluster cells by PhenoGraph using the Leiden algorithm (*k* = 30, and resolution parameter = 0.5). Differential gene expressions per cluster were computed using the Wilcoxon test by comparing each cluster to the rest of clusters. For visualization, we imputed the expression of genes using MAGIC ^79^ (nPCA = 30, *knn* = 30). Imputed data were ensured to be consistent with the original non-imputed data.

### *In situ* proximity ligation assay

Endogenous protein interactions were detected using an *in situ* PLA kit according to the manufacturer’s instruction (Sigma Aldrich Cat# DUO92101). Briefly, cells were seeded on a chamber slide for 24 h, fixed in 4% PFA for 10 min at room temperature, and permeabilized with 0.01% Triton-X in PBS for 10 min at room temperature. Cells were then incubated with Duolink blocking solution for 1 h at 37°C and two primary antibodies overnight at 4°C. Antibodies used were rabbit anti-CHD1 (Invitrogen, Cat# PA5-82795) and mouse anti-c-Jun (Invitrogen Cat# MA5-15881) or rabbit anti-CELSR1 (Millipore Cat# ABT119) and mouse anti-L1CAM (Invitrogen Cat# MA5-14140). Samples were washed twice with the manufacturer’s Wash Buffer A at room temperature, and were incubated with Duolink *In Situ* PLA anti-mouse Plus and anti-rabbit Minus probes for 1 h at 37°C. After two times washing with Wash Buffer A, samples were incubated with the manufacturer’s ligase for 30 min at 37°C. Oligonucleotides conjugated to the probes were amplified using the proprietary polymerase provided by the manufacturer for 100 min at 37°C. The samples were washed two times with the manufacturer’s Wash Buffer B and one time with 0.01x Wash Buffer B, and were mounted to a coverslip with ProLong diamond antifade mountant (Life Technologies Cat# P36970) before microscopy.

### Flow cytometry

Analysis and sorting were conducted on a FACS Aria II SORP flow cytometer (Beckton Dickinson) or FACSymphony S6 high-speed cell sorter (Beckton Dickinson) using a 100 μm nozzle. Cells were resuspended in cold PBS supplemented with 2% FBS, filtered through a 40-μm cell strainer, and counterstained with DAPI (1 μg/mL). When sorting for L1CAM^+^ KP cells, single-cell samples were incubated with antibody against mouse L1CAM conjugated with allophycocyanin (Miltenyi Biotec Cat# 130-102-221) for 1 h in ice and washed three times with cold PBS with 2% FBS. Cells were gated by forward and orthogonal light scatter before analyzing with 405, 488, 561 and/or 640 nm excitation depending on experimental parameters. Fluorescence signals were measured with logarithmic amplifiers, and cells of interest were sorted according to the spectral separation between the samples and controls. Sorted cells were either proceeded directly for single-cell sequencing analysis or placed in tissue culture for experiments.

### Bioinformatics analysis

For pathway analysis, a differentially enriched gene list was derived by comparing the L1CAM^+^SOX2^+^ cluster #12 versus the rest of clusters from the scRNA-seq day 7 tumoroid dataset. The 3000 top scoring genes were curated and processed to identify uniquely enriched biological pathways based on the Gene Ontology (GO), Reactome, and PANTHER databases. Candidate pathways were cross-referenced across the databases, and pathways shared by all three databases were further analyzed for component biological processes. The processes were ranked based on the accumulated negative log-transformed false discover rate (FDR), and pathways were evaluated based on their enrichment scores.

To identify transcription regulators associated with *Sox2* expression, the differentially enriched gene list from the same L1CAM^+^SOX2^+^ scRNA-seq cluster was analyzed by Binding Analysis for Regulation of Transcription (BART). Ranked transcription regulators, including transcription factors and epigenetic modifiers, were identified by relative scores. Based on ranked scores, the top 50 candidates were selected and additionally filtered based on the ChIP Enrichment Analysis (CHEA) and Encyclopedia of DNA Elements (ENCODE) databases for involvement in *Sox2* expression.

### Statistical analysis

Sample sizes were indicated in figures or figure legends. An unpaired two-tailed Mann–Whitney test, two-tailed t test or two-tailed Fisher’s exact test was used for two-group comparisons. If multiple groups were compared, one-way analysis of variance was performed, followed by the Tukey test. Tumor BLI curves were tested using Kolmogorov-Smirnov test, and survival curves were tested using log-rank (Mantel-Cox) test. Data normality was tested using the Shaprio-Wilk normality test. All statistical analyses were performed using GraphPad Prism v.10.2.3 for Mac (GraphPad Software, www.graphpad.com). All bar graphs show mean values with error bars representing either s.e.m. or s.d., as defined in legends. *P* values are indicated in the figure legends and, when appropriate, were rounded to the nearest single significant digit. *P* values of less than 1 × 10^−4^ were indicated as a range. *P* values of less than 0.05 were considered to be significant.

## Data availability

The sequencing data have been submitted to the Gene Expression Omnibus under accession number GSE284443.

---

**Extended Data Figure 1.**
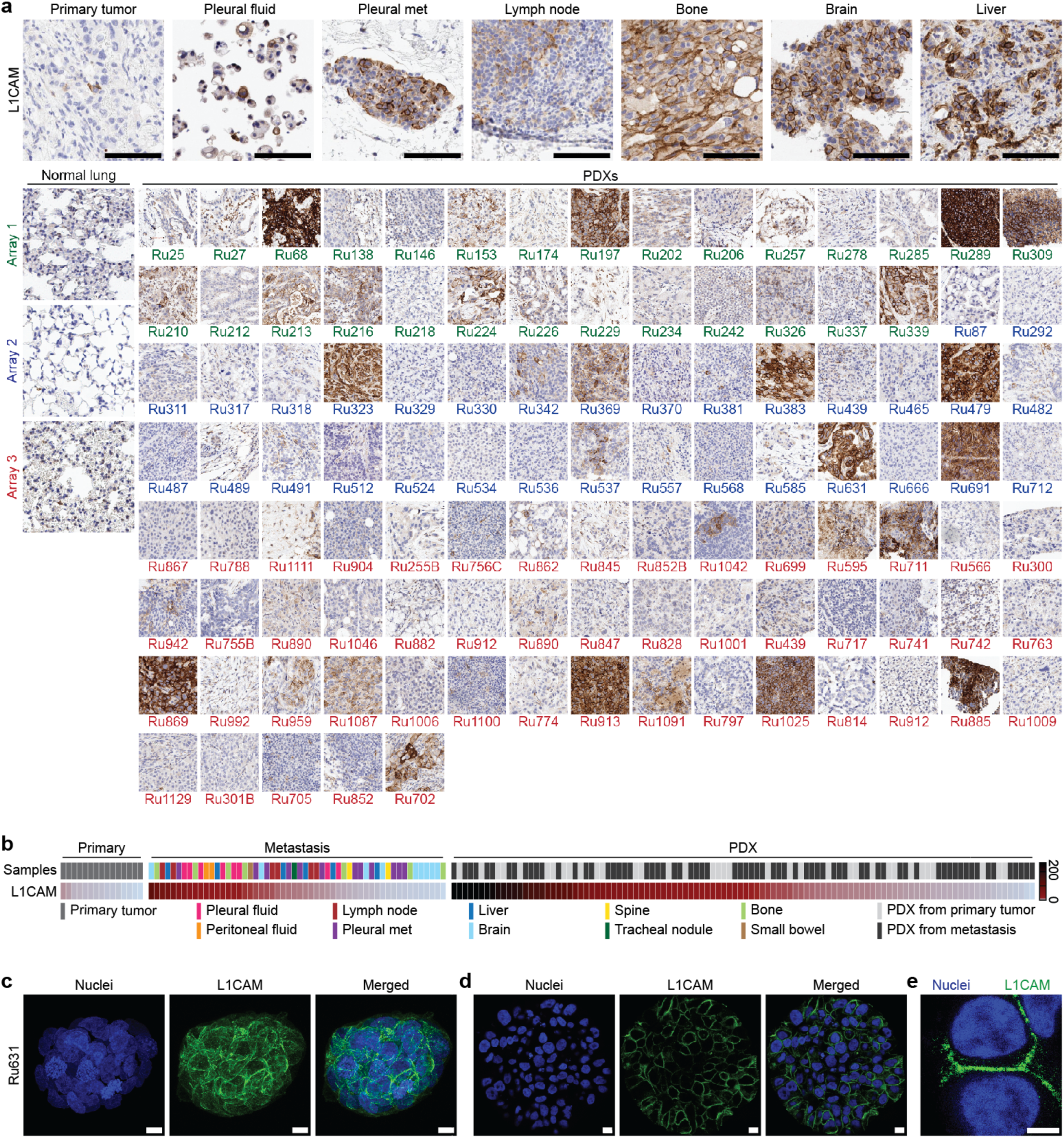
L1CAM expression in patient tissues, PDXs, and PDX tumoroids. **a**, Representative L1CAM IHC staining in primary tumors, pleural fluids, and metastatic samples from LUAD patients and three tissue microarrays (Array 1, green; Array 2, blue; Array 3, red) containing normal human lung tissue and LUAD PDX (Passage 0 or 1). Scale bar, 100 μm. **b**, Heatmap of L1CAM IHC staining quantification (H-score) in primary tumor, pleural fluid, metastatic samples, and PDXs from LUAD patients. Each column represents an individual sample. Samples are color-coded according to their site of tissue. **c**, Maximum projection of a PDX tumoroid fluorescently stained with L1CAM. Scale bar, 10 μm. **d**, Cross-section of a PDX tumoroid stained with L1CAM IF and DAPI (nuclei). Scale bar, 10 μm. **e**, Close up of a PDX tumoroid stained with L1CAM IF and DAPI. Scale bar, 5 μm.

**Extended Data Figure 2.**
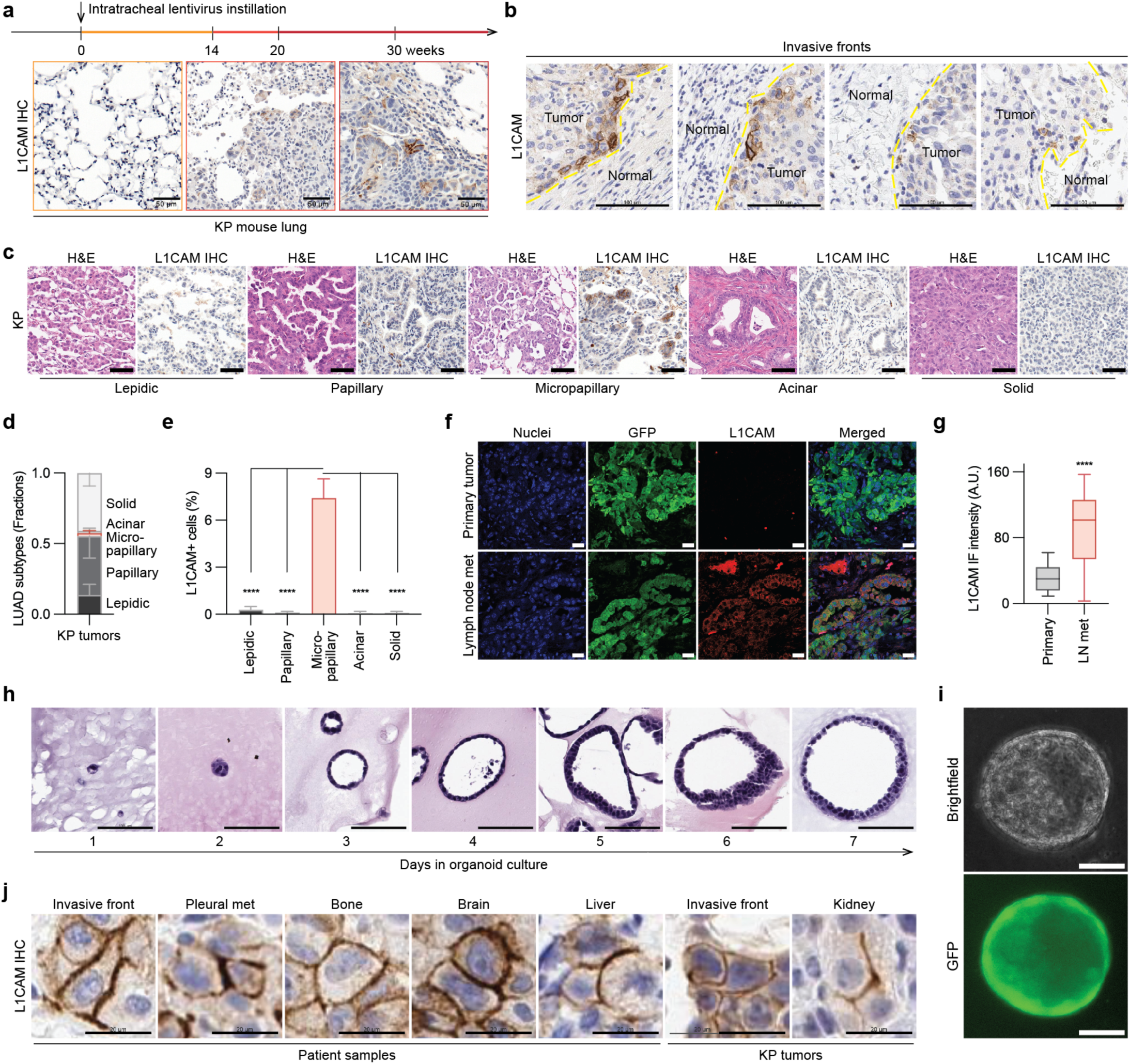
L1CAM expression in KP mouse tissues and KP tumoroids. **a**, L1CAM IHC staining in lung and tumor sections in KP mice at baseline and at different time points after lentivirus instillation to induce adenocarcinoma formation. Yellow, week 0; orange, week 14; red, week 24. Scale bar, 50 μm. **b**, L1CAM IHC staining of the patient tissue sections including normal tissue and tumor. The invasive tumor front is indicated with a yellow line. Scale bar, 200 μm. **c**, H&E and L1CAM IHC staining of KP primary tumor sections (26 week post-Cre) showing the indicated histological subtypes. Scale bar, 50 μm. **d**, Fraction of histological subtypes observed in KP primary tumors. *n* = 203,870 cells. **e**, Percentage of L1CAM^+^ cells in KP primary tumors with respect to histological subtypes. Lepidic (*n =* 26,871); Papillary (*n =* 88,381); Micropapillary (red colored, *n =* 4,116); Acinar (*n =* 3,430); Solid (*n =* 81,072). \*\*\*\**P* < 0.0001. **f**, Representative GFP (cancer cells) IF and L1CAM IF images of primary tumor and an autochthonous lymph node metastasis from a KP mouse (35 week post-Cre). Scale bar, 20 μm. **g**, Quantification of L1CAM IF intensity in the experiment (**f**). LN, lymph node; AU, arbitrary unit. *n =* 24 image frames; *N* = 3 mice in each condition. \*\*\*\**P* < 0.0001. **h**, H&E staining of KP tumoroids grown in Matrigel over the course of 7 days. Scale bar, 100 μm. **I**, Brightfield and GFP fluorescence microscopy images of KP tumoroids at day 7. Scale bar, 10 μm. **j**, Close up of L1CAM IHC staining of patient samples and KP tumors. Scale bar, 20 μm. Statistical significance was assessed using the two-tailed Mann-Whitney test (**g**) or one-way analysis of variance followed by the Tukey test (**e**). Data are shown as mean ± S.D. (**d**,**e**).

**Extended Data Figure 3.**
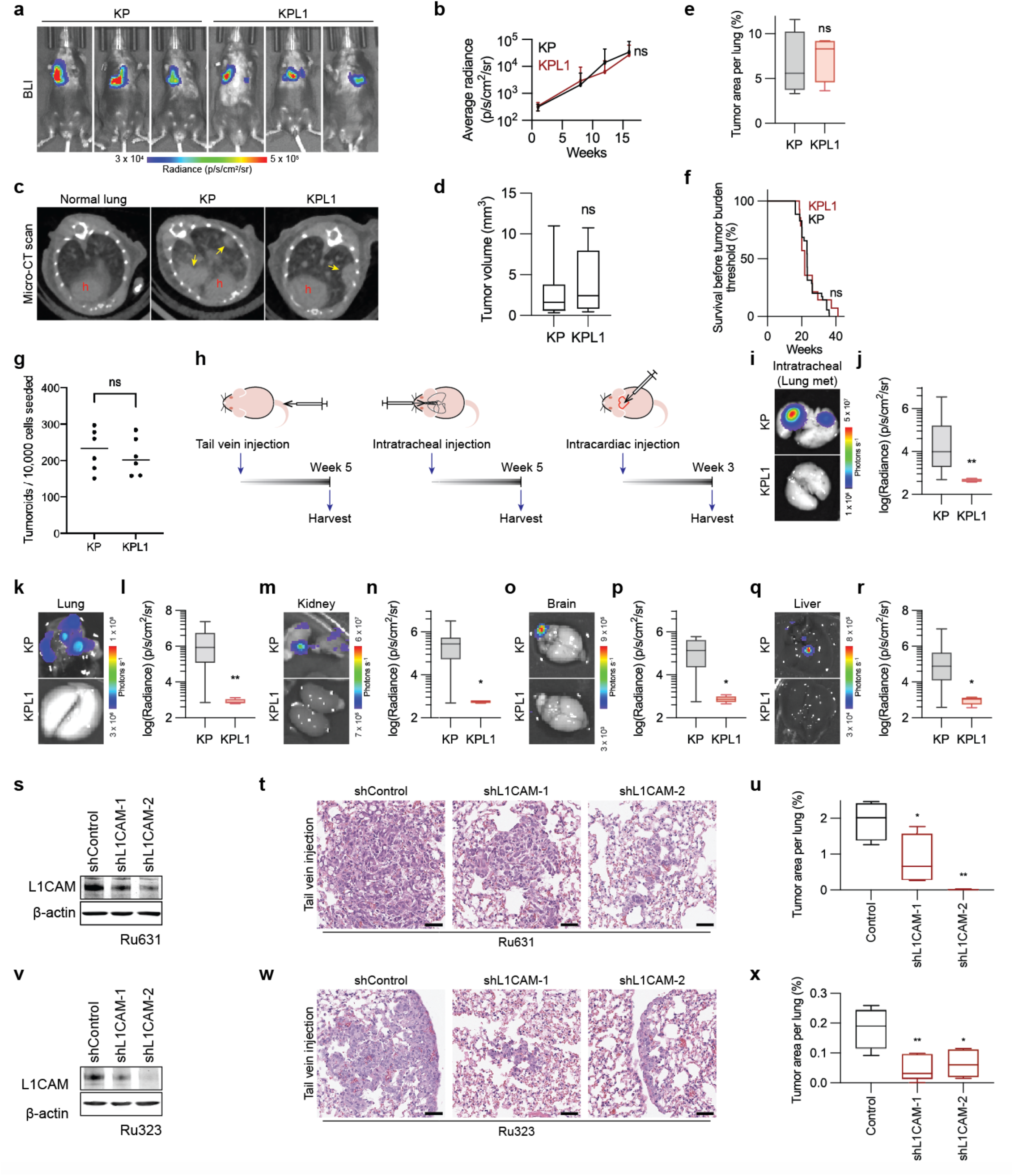
L1CAM promotes LUAD metastasis but not tumor initiation. **a**, Representative images of tumor-bearing KP and KPL1 mice analyzed by BLI at week 16 after viral transduction. **b**, Growth curve of KP and KPL1 primary tumor burden as determined by BLI. KP, *n =* 37; KPL1, *n =* 14. ns, *P* > 0.9999. **c**, Micro-CT scan images of normal lung and tumor-bearing lungs from KP and KPL1 mice at week 22 after viral transduction. *Arrows*, tumors; *red h*, heart. **d**, Tumor burden comparison between KP and KPL1 mice based on micro-CT scans at week 22. *n =* 38 KP mice, *n =* 10 KPL1 mice. ns, *P* = 0.3192. **e**, Tumor areas of KP and KPL1 mouse lungs at week 22-30 after viral transduction. *n* = 4 mice. ns, *P* = 0.6857. **f**, Graphical representation of mouse survival prior to reaching the target tumor burden, set at a luciferase bioluminescence flux of 10^7^ photons/sec per mouse chest. KP, *n* = 35; KPL1, *n* = 14. ns, *P* = 0.7143. **g**, Quantification of tumoroids formed per 10,000 cells over 7 days. KP, *n =* 6; KPL1, *n =* 6. ns, *P* = 0.6991. **h**, Schematic of three different metastasis inoculation routes. **i**, Representative image of *ex vivo* lung BLI signals at week 5 after intratracheal delivery of single-cell suspension of KP and KPL1 tumoroids (2 x 10^4^ cells) into athymic mice. **j**, Quantification of *ex vivo* lung BLI signal in the experiment in panel (**i**). KP, *n =* 8; KPL1, *n =* 6. \*\**P* = 0.0027. **k**, Representative image of *ex vivo* lung BLI signals at week 3 after intracardiac injection of single-cell suspension of KP and KPL1 tumoroids (5 x 10^4^ cells) into athymic mice. **l**, Quantification of *ex vivo* lung BLI signals in the experiment in panel (**k**). KP, *n =* 9; KPL1, *n =* 5. \*\**P* = 0.0070. **m**, Representative image of *ex vivo* kidney BLI signals at week 3 after intracardiac injection of single-cell suspension of KP tumoroids or KPL1 tumoroids (5 x 10^4^ cells). **n**, Quantification of *ex vivo* kidney BLI signals at week 3 after intracardiac injection of single-cell suspension of KP tumoroids or KPL1 tumoroids. KP, *n =* 9; KPL1, *n =* 5. \**P* = 0.0120. **o**, Representative image of *ex vivo* brain BLI signals at week 3 after intracardiac injection of single-cell suspension of KP tumoroids or KPL1 tumoroids (5 x 10^4^ cells). **p**, Quantification of *ex vivo* brain BLI signals at week 3 after intracardiac injection of single-cell suspension of KP tumoroids or KPL1 tumoroids. KP, *n =* 9; KPL1, *n =* 5. \**P* = 0.0120. **q**, Representative image of *ex vivo* liver BLI signals at week 3 after intracardiac injection of single-cell suspension of KP tumoroids or KPL1 tumoroids (5 x 10^4^ cells). **r**, Quantification of *ex vivo* liver BLI signals at week 3 after intracardiac injection of single-cell suspension of KP tumoroids or KPL1 tumoroids. KP, *n =* 9; KPL1, *n =* 5. \**P* = 0.0120. **s**, Western immunoblotting analysis of Ru631 PDX cells transduced with two different *L1CAM* shRNAs. **t**, H&E staining of lung sections four weeks after injecting Ru631 L1CAM knockdown via tail vein. Scale bar, 50 μm. **u**, Quantification of Ru631 tumors after tail vein injection. *n* = 4 mice per condition. \**P* = 0.0337; \*\**P* = 0.0013. **v**, Western immunoblotting of L1CAM knockdowns with two different short hairpins in Ru323 PDXs. **w**, H&E staining of lung sections four weeks after injecting Ru323 with L1CAM knockdown via tail vein. Scale bar, 50 μm. **x**, Quantification of Ru323 tumors after tail vein injection. *n* = 5 mice per condition. *\*P* = 0.0123; \*\**P* = 0.0058. Data are shown as a box (median ± 25-75%) and whisker (maximum to minimum values) plot (**d**,**e**,**j**,**l**,**n**,**p**,**r**). Error bar indicates mean ± S.D. (**b**). Statistical significance was assessed using Kolmogorov-Smirnov test (**b**), two-tailed Mann-Whitney test (**d**,**e**,**g**,**j**,**l**,**n**,**p**,**r**) or one-way analysis of variance followed by the Tukey test (**u**,**x**).

**Extended Data Figure 4.**
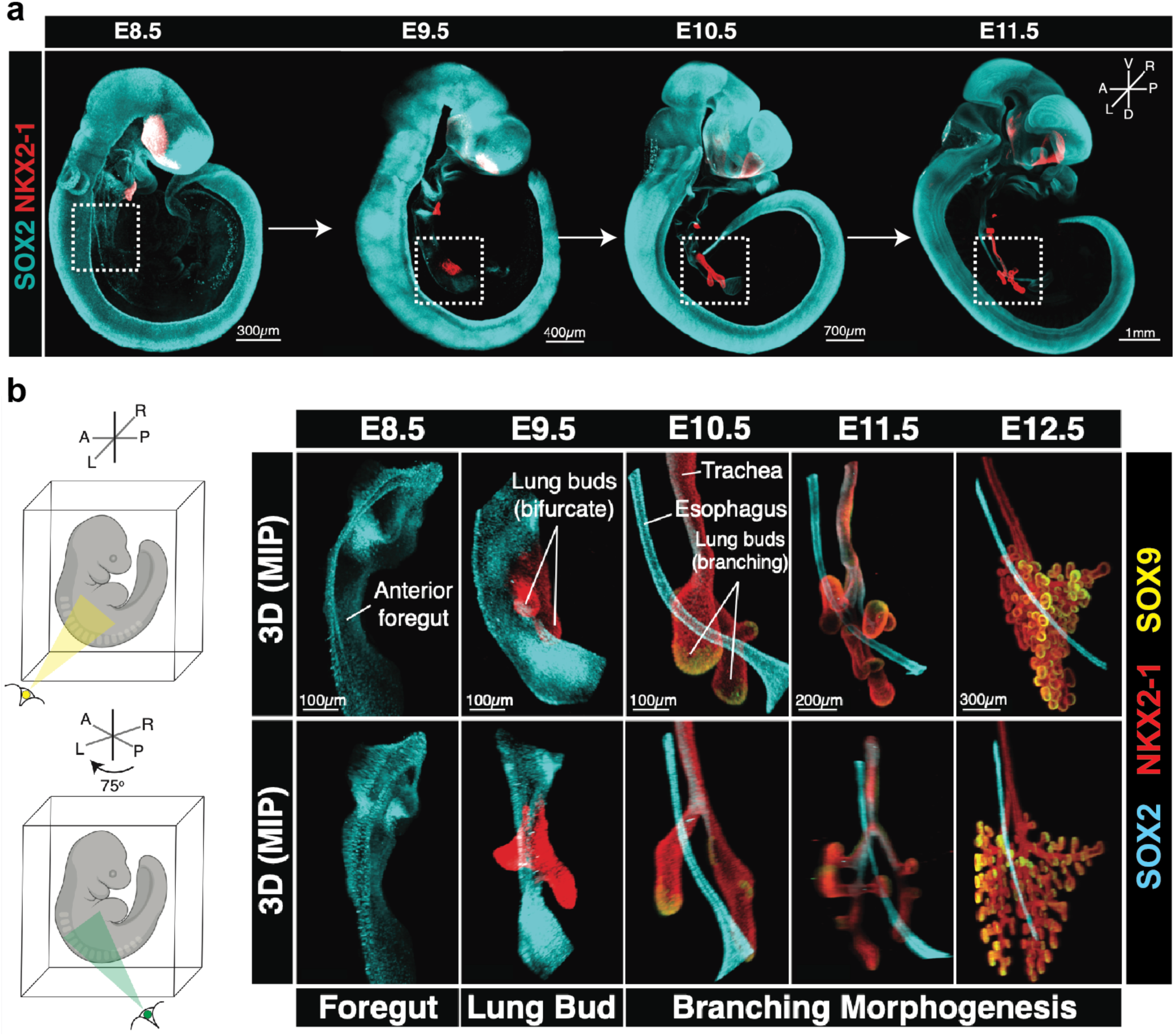
Lung developmental continuum in mouse embryo. **a**, Representative iDISCO-cleared whole-mount 3D rendered Maximum Intensity Projections (MIP) images of sequentially staged wild type mouse embryos stained by IF for SOX2 and NKX2-1 from E9.5 to E11.5. The dashed box demarcates a portion of the developing respiratory tract depicted in higher resolution in subsequent panels. Scale bars are defined in the panels. **b,** Representative 3D MIP images of SOX2, NKX2-1 and SOX9 IF staining during mouse lung development. The bottom panels are 75° rotated views of the respiratory region. All data were independently validated from replicate samples of at least *n =* 4 embryos with similar results obtained. *E*, embryonic day; *A*, anterior; *P*, posterior; *L*, left; *R*, right. Scale bars are defined in the panels.

**Extended Data Figure 5.**
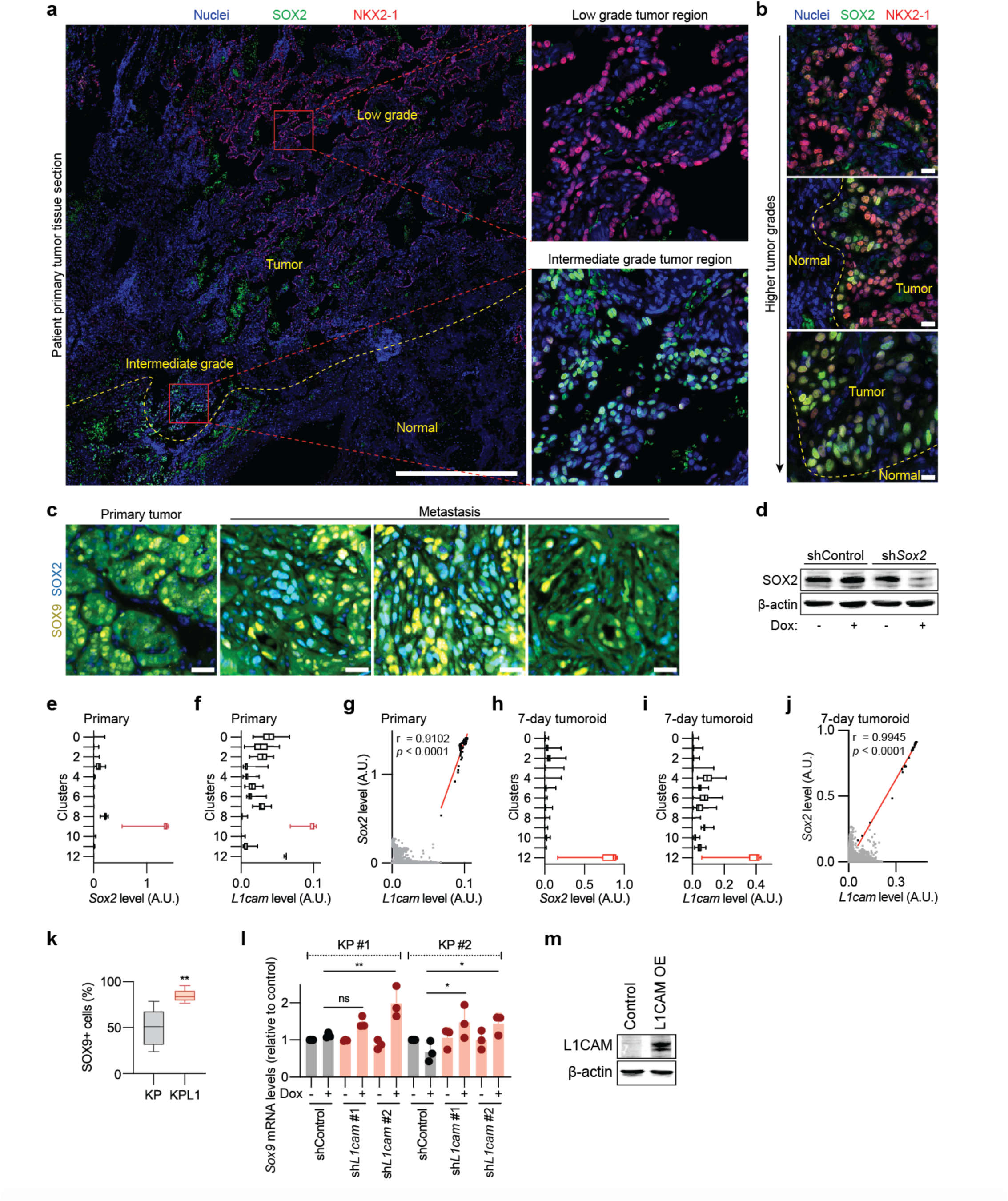
L1CAM-dependent LUAD metastasis linked to SOX2 expression. **a**, SOX2 and NKX2-1 IF staining of patient-derived LUAD primary tumor tissue section. The invasive front is marked with a dotted yellow line, and the magnified low-grade and high-grade tumor areas are shown in red boxes. Scale bar, 1 mm. **b**, SOX2 and NKX2-1 IF staining of patient-derived primary tumor tissue section with different tumor grade areas. Invasive front is shown with a dotted yellow line. Scale bar, 20 μm. **c**, SOX2 and SOX9 IF staining in KP primary tumor and metastasis tissue sections collected simultaneously from the same mouse (35 week post-Cre). Scale bar, 100 μm. **d**, Western immunoblotting analysis of SOX2 expression after dox-induced *Sox2* knockdown in KP tumoroids. **e**,**f**, Box and whisker plots of the imputed transcript levels of *Sox2* (**e**) and *L1cam* (**f**) in the scRNA-seq transcriptional clusters from KP primary tumors. **g**, Scatter plot of *L1cam* versus *Sox2* expression in the scRNA-seq dataset from KP primary tumors. Data from transcriptional cluster 9 are shown in black. **H**,**I**, Box and whisker plots of the imputed transcript levels of *Sox2* (**h**) and *L1cam* (**i**) from 7-day tumoroids. Data from the cluster 12 are shown in red. **j**, Scatter plot of *L1cam* versus *Sox2* expression in the scRNA-seq dataset from 7-day tumoroids. Data from transcriptional cluster 12 are shown in black. **k**, Proportion of SOX9^+^ cells detected by IF in lung colonies after injecting a high number (1 x 10^5^) of KP or KPL1 cells via the tail vein into athymic mice. KP, *n =* 10; KPL1, *n =* 5. \*\**P* = 0.0013. **l**, Relative expression of SOX9 in two independent KP tumoroid lines upon conditional knockdown of L1CAM. Bar graphs, mean ± S.D. *n* = 3 in each condition. ns, *P* = 0.8301; \**P* = 0.0202, 0.0305; \*\**P* = 0.0094. **m**, Western immunoblotting analysis of L1CAM levels after L1CAM overexpression in KPL1 tumoroids. Data are shown as a box (median ± 25-75%) and whisker (maximum to minimum values) plot (**e**,**f**,**h**,**i**,**k**). Spearman correlation was used to calculate the relationship between L1CAM expression and SOX2 expression (**g**,**j**). Statistical significance was assessed using the two-tailed Mann-Whitney test (**k**) or one-way analysis of variance followed by the Tukey test (**l**).

**Extended Data Figure 6.**
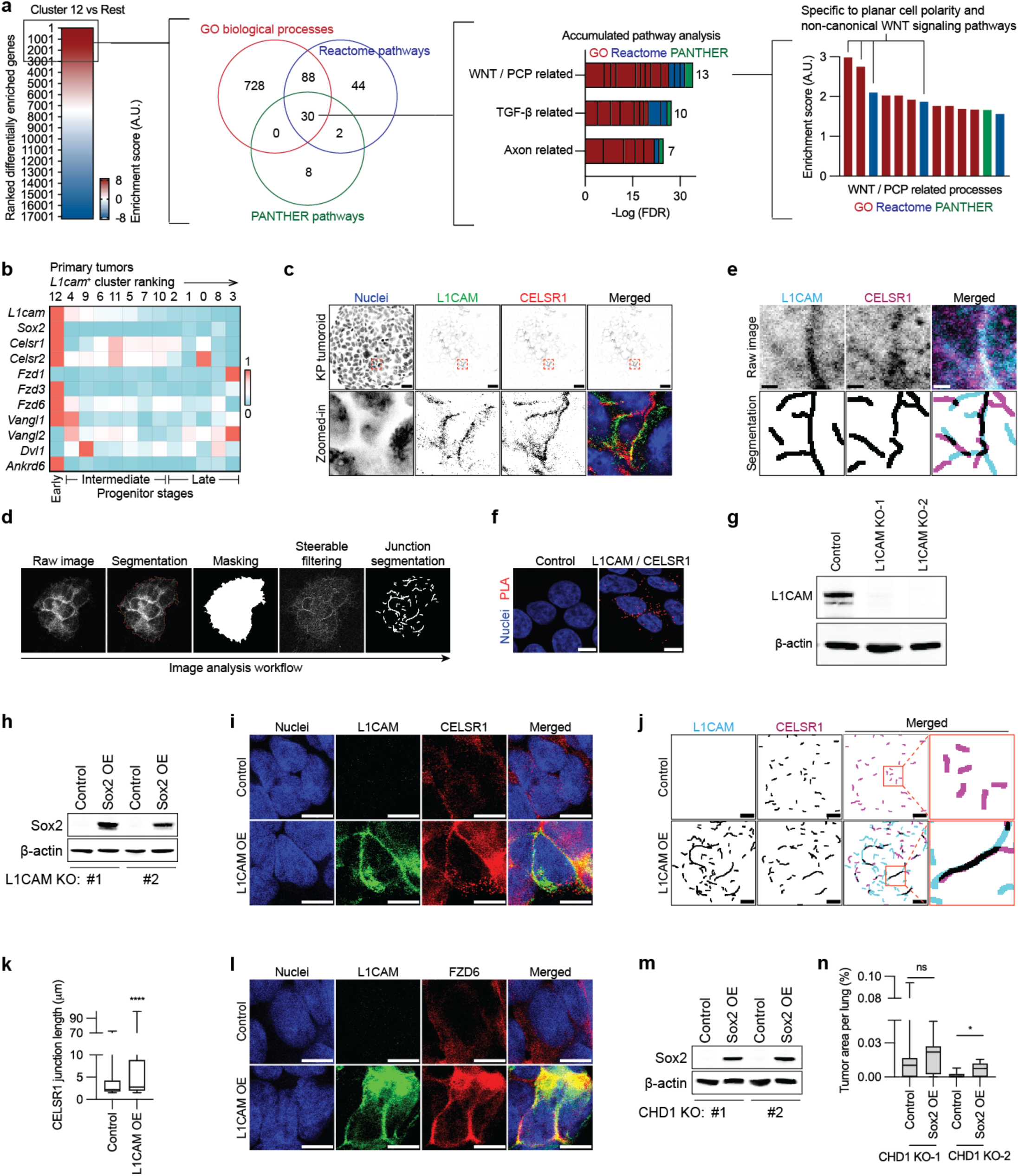
Analysis of PCP components in LUAD L1CAM^+^/SOX2^+^ cells. **a**, WNT/PCP pathway scoring as the top hit among highly enriched genes in L1CAM^+^/SOX2^+^ KP tumoroid cells. Top 3000 ranked differentially expressed genes in cluster #12 versus the rest of cell population in day-7 KP tumoroids were analyzed to determine uniquely enriched biological pathways from the Gene Ontology (GO), Reactome, and PANTHER databases. 30 pathways shared among these three analyses were concatenated into three related processes. 13 of these pathways were from the WNT/PCP-related process and had the lowest accumulated false discovery rate (FDR). Within the WNT/PCP-related processes, the PCP and non-canonical WNT signaling pathways showed the highest enrichment score. **b**, Heatmap displaying the average expression of core PCP components in the primary tumor clusters defined in Figure 2L. The clusters are ranked left to right according to the average *L1cam* expression level. **c**, L1CAM and CELSR1 IF staining in KP tumoroids. The magnified region is indicated by *red* or *white boxes*. Scale bar, 10 μm. **d**, Image analysis workflow for detecting the colocalization of L1CAM and CELSR1 at cell-cell junctions using a steerable filter to extract curvilinear image features. **e**, Computer vision-driven detection and segmentation of L1CAM and CELSR1 at cell-cell junctions through steerable filtering. The thickness of segmented junctions was dilated for visualization purposes. Scale bar, 1 μm. **f**, L1CAM and CELSR1 PLA fluorescence in H23 LUAD cells. The PLA signals are shown as red dots with nuclei counterstaining. Scale bar, 10 μm. **g**, L1CAM western immunoblotting analysis in parental and L1CAM knockout H23 cells. **h**, Western immunoblotting analysis of SOX2 overexpression in H23 LUAD cells upon CRISPR/Cas9-mediated *L1CAM* knockout, clones 1 and 2. **i.** L1CAM and CELSR1 IF staining in HEK293T cells engineered to overexpress L1CAM versus control. Scale bar, 10 μm. **j**, L1CAM and CELSR1 segmentation analysis at cell-cell junctions in L1CAM overexpressing HEK293T cells. The region of higher magnification is indicated by a *red box*. **k**, Quantification of CELSR1 at cell-cell junctions in control or L1CAM overexpressing HEK293T cells. Control, *n =* 3406 junctions; L1CAM OE, *n =* 6508 junctions. \*\*\*\**P* < 0.0001. **l**, L1CAM and FZD6 IF staining in control or L1CAM overexpressing HEK293T cells. Scale bar, 10 μm. **m**, Western immunoblotting analysis of SOX2 overexpression in CHD1 knockout H23 cells. **n**, Box and whisker plots showing the size of metastatic lesions per lung in SOX2 overexpressing, CHD1 knockout H23 cells. *n =* 9 for CHD1 KO-1; 10 for CHD1 KO-2. ns, *P* = 0.2805; \**P* = 0.0415. Data are shown as a box (median ± 25-75%) and whisker (maximum to minimum values) plot (**k**,**n**). Statistical significance was assessed using the one-way analysis of variance followed by the Tukey test (**k**) or two-tailed Mann-Whitney test (**n**).

